# Tailoring evidence into action: using a codesign approach for biodiversity information in the Tropical Andes

**DOI:** 10.1101/2023.05.20.541564

**Authors:** Jose W. Valdez, Henrique M. Pereira, Gustavo Francisco Morejón, Cristina Acosta-Muñoz, Francisco Javier Bonet Garcia, Lucía Castro Vergara, Claros R. Xavier, Michael J. Gill, Carmen Josse, Indyra Lafuente-Cartagena, Robert Langstroth, Sidney Novoa Sheppard, Gabriela Orihuela, Francisco J. Prieto-Albuja, Natividad Quillahuaman, Marcos F. Terán, Carlos M. Zambrana-Torrelio, Laetitia M. Navarro, Miguel Fernandez

**Affiliations:** German Centre for Integrative Biodiversity Research (iDiv) Halle-Jena-Leipzig, Puschstrasse 4, 04103, Leipzig, Germany; Institute of Biology, Martin Luther University Halle Wittenberg, Am Kirchtor 1, 06108 Halle (Saale), Germany; Estación Biológica de Doñana Departamento de Biología de la Conservación Américo Vespucio n° 26 – 41092 Sevilla, Spain; Fundación EcoCiencia, San Ignacio E12-143 y Humboldt. Edificio Carmen Lucia Dept. 1 (Sector González Suárez - Norte de Quito). CP: 170517 Quito-Ecuador; Department of Botany, Ecology and Plant Physiology, University of Cordoba. C.U. Rabanales, 14014 Cordoba, Spain; Asociación para la Conservación de la Cuenca Amazónica - ACCA, Calle Vargas Machuca 627, Miraflores, Lima, Peru; Asociación para la Investigación y Conservación de Ecosistemas Andino Amazónicos (ACEAA-Conservación Amazónica), Bolivia; NatureServe, 2550 South Clark St. Suite 930, Arlington VA, 22202, U.S; 43611 Hetrick Ln, South Riding, VA 20152, USA; Instituto Nacional de Biodiversidad INABIO. Pasaje Rumipamba 341 y Av. de los Shyris. Código Postal 170506. Quito-Ecuador; Department of Environmental Science and Policy, George Mason University, 4400 University Dr. Fairfax VA, 22030, U.S; Instituto de Ecología, Universidad Mayor de San Andrés, Calle 27 Cota-cota, La Paz, Bolivia

**Keywords:** stakeholder engagement, biodiversity monitoring, essential biodiversity variables, Ecuador, Peru, policy, Bolivia, monitoring, mainstreaming

## Abstract

Biodiversity conservation is a complex and transdisciplinary problem that requires engagement and cooperation among scientific, societal, economic, and political institutions. However, historical approaches have often failed to bring together and address the needs of relevant stakeholders in decision-making processes. The Tropical Andes, a biodiversity hotspot where conservation efforts often conflict with socioeconomic issues and policies that prioritize economic development, provides an ideal model to develop and implement more effective approaches. In this study, we present a codesign approach that mainstreams and improves the flow of biodiversity information in the Tropical Andes, while creating tailored outputs that meet the needs of economic and societal stakeholders. We employed a consultative process that brought together biodiversity information users and producers at the local, national, and regional levels through a combination of surveys and workshops. This approach identified priority needs and limitations of the flow of biodiversity information in the region, which led to the co-design of user-relevant biodiversity indicators. By leveraging the existing capacities of biodiversity information users and producers, we were able to co-design multiple biodiversity indicators and prioritize two for full implementation ensuring that the data was findable, accessible, interoperable, and reusable based on the FAIR principles. This approach helped address limitations that were identified in the stakeholder engagement process, including gaps in data availability and the need for more accessible biodiversity information. Additionally, capacity-building workshops were incorporated for all stakeholders involved, which aimed to not only improve the current flow of biodiversity information in the region but also facilitate its future sustainability. Our approach can serve as a valuable blueprint for mainstreaming biodiversity information and making it more inclusive in the future, especially considering the diverse worldviews, values, and knowledge systems between science, policy, and practice.

## Introduction

The Tropical Andes is a biodiversity hotspot where conservation efforts collide with socioeconomic issues and public policies prioritizing economic development (Fernández et al. 2015; Josse & Fernandez 2021; Rodríguez-Echeverry & Leiton 2021). Despite covering less than half a percent of the Earth’s surface, this region contains over 10 percent of globally described species across 100 distinct ecosystems that provide vital provisioning, cultural, and regulating services to several South American countries (Myers et al. 2000; Rodríguez-Mahecha et al. 2004; Anderson et al. 2011; Josse et al. 2011). However, deforestation, mining, and other unsustainable practices threaten the region’s biodiversity and the well-being of its inhabitants, supported by significant investments from multilateral financial organizations (Jetz et al. 2007; Jarvis et al. 2010; Armenteras et al. 2011; Josse et al. 2011; Rodríguez et al. 2013; Romero-Muñoz et al. 2019; Rodríguez-Echeverry & Leiton 2021). Protecting the biodiversity and ecosystems of the Tropical Andes is essential for both the world’s species and the well-being of millions who depend on its ecosystems.

Most efforts to halt and reverse biodiversity declines typically involve conservation policies that aim to balance protection, restoration, and sustainable use with societal and economic development (Smith et al. 2020); yet their failure highlights a gap between the scientific community, society, businesses, and policymakers (Diedrich et al. 2011; Jolibert & Wesselink 2012; Fernández-Llamazares & Rocha 2015; Smith et al. 2020; World Economic Forum 2021; Xu et al. 2021). Biodiversity conservation is a complex, multi-causal problem that requires engagement and cooperation across scientific, societal, economic, and political institutions to meet the needs of all stakeholders (Jolibert & Wesselink 2012). However, historically, input from relevant sectors has been lacking (Dempsey 2013; Neßhöver et al. 2013; Pisupati & Prip 2015; Karlsson-Vinkhuyzen et al. 2017). Despite all groups of society being vulnerable to biodiversity loss to varying degrees, economic, development, and societal sectors have typically been considered incompatible with conservation goals, leading to a perception that such goals do not align with their interests (Folke 2006; Smith et al. 2020; Morley et al. 2021).

To identify effective conservation actions and foster engagement and ownership across stakeholders, cooperation and communication among diverse interest groups are essential (Pascual et al. 2021; Perino et al. 2021). Achieving this requires a collaborative, cross-sectoral, and multinational approach that takes a pluralistic perspective on biodiversity given the multiple worldviews, values, and knowledge systems between science, policy, and practice (Zador et al. 2015; Bravo et al. 2016; Pascual et al. 2021; Mansur et al. 2022). One promising approach is ‘biodiversity mainstreaming’ whereby biodiversity and conservation considerations are integrated and embedded into the strategies and policies of key economic and societal sectors that impact or rely on biodiversity (Chandra & Idrisova 2011; Huntley 2014; Redford et al. 2015; Whitehorn et al. 2019). Biodiversity mainstreaming has already gained significant traction and has been incorporated by the United Nations Convention on Biological Diversity (CBD), International Union for Conservation of Nature (IUCN), European Union (EU) Biodiversity Strategy, National Biodiversity Strategies and Action Plans (NBSAPs), The Intergovernmental Platform on Biodiversity and Ecosystem Services (IPBES), and global efforts such as the post-2020 Global Biodiversity Framework (Huntley 2014; Josse & Fernandez 2021; Perino et al. 2021). Despite the widespread application of the biodiversity mainstreaming process, there is currently a lack of established guidelines, recognized best practices, and empirical evidence on its effectiveness, which limits its integration into decision-making processes and makes its impacts unclear (Huntley 2014).

Due to the complex and multifaceted nature of mainstreaming biodiversity, there are often numerous limitations and bottlenecks in the flow of biodiversity information from producers to users (data collection, analysis, decision-making, and dissemination; Figure 1), which can hinder the effective implementation and evaluation of mainstreaming initiatives. One issue is the lack of accessible and standardized data across different institutions, sectors, and regions, which impedes informed decision-making and assessment of the impacts of actions on biodiversity (Stephenson et al. 2017). This problem is further compounded by the lack of coordination and communication among stakeholders across different levels of data flow, as well as the absence of clear policies and guidelines for mainstreaming biodiversity into decision-making (Navarro et al. 2017; Josse & Fernandez 2021). This often leads to confusion, conflicting information, and the risk of duplicating efforts and investments. Capacity limitations among stakeholders due to a lack of resources or expertise, particularly in under-developed regions such as the Tropical Andes, also contribute to bottlenecks within the flow of biodiversity information (Fernández et al. 2015; Josse & Fernandez 2021; Alvarado et al. 2022). Additionally, the challenges and limitations of biodiversity mainstreaming vary at different scales, making it difficult to generalize solutions across local, national, and regional contexts (Karlsson-Vinkhuyzen et al. 2014; Alvarado et al. 2022).

**Figure 1.**
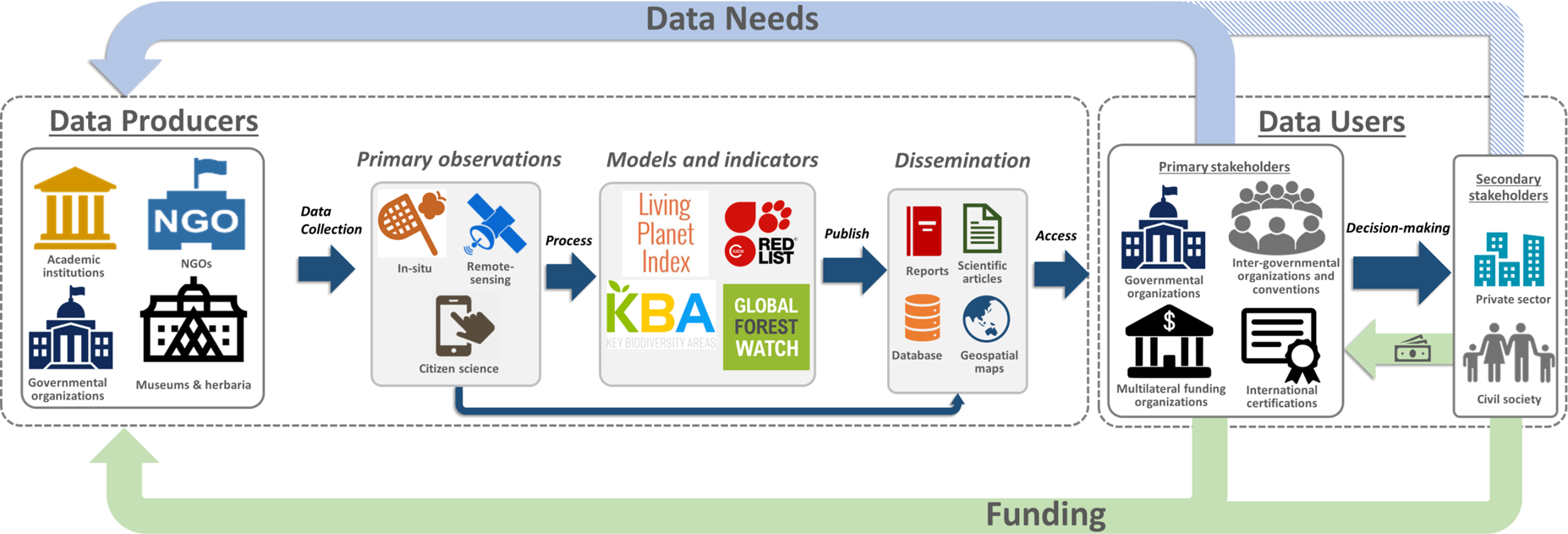
Mainstreaming the flow of biodiversity information between producers and users. Primary stakeholders are those groups and individuals who directly pay for or ask for biodiversity information. Secondary stakeholders are typically not directly involved in the process of information flow. The flow of information between producers and users relies on the resources (funding and capacities) of the data producers.

Another main challenge of biodiversity mainstreaming is the lack of engagement with relevant stakeholders and the disconnect between producers of biodiversity data and potential end-users (Figure 1). The primary stakeholders that typically use biodiversity information are those who directly request and/or pay for it, and sometimes even collect it, such as policymakers, conservation practitioners, land managers, multilateral funding organizations, NGOs, inter-governmental organizations and conventions, and researchers (Figure 1). These primary stakeholders rely on up-to-date and accurate information to make informed decisions regarding biodiversity and conservation management. Another group, which we refer to as “secondary stakeholders”, such as businesses, civil society groups, local communities, and the general public (Figure 1), also benefit from and attributes values to biodiversity, but are not typically involved in the mainstreaming process due to limited resources, perceived lack of knowledge or interest, priority towards primary stakeholders, and limited recognition of their perspectives and contributions (Jolibert & Wesselink 2012; Neßhöver et al. 2013; Smith et al. 2020). However, since secondary stakeholders have the potential to significantly influence policies and funding decisions that affect biodiversity, involving them in the mainstreaming process can also help raise awareness of the value and importance of biodiversity (Josse & Fernandez 2021; Alvarado et al. 2022). Until now, barriers such as communication gaps, a narrow focus on environmental benefits, and a government and academic-driven approach often leave stakeholders feeling ignored and contribute to power imbalances, further hindering mainstreaming efforts (Vogel et al. 2007; Chandra & Idrisova 2011; Cvitanovic et al. 2016; Josse & Fernandez 2021; Alvarado et al. 2022).

To address the challenges of biodiversity mainstreaming and to improve the flow of biodiversity information, it is crucial to implement strategies that involve all relevant sectors and groups (Sterling et al. 2017; Gavin et al. 2018; Alvarado et al. 2022). This includes participatory research, multi-stakeholder dialogues, and adaptive frameworks, which can create a more comprehensive and inclusive mainstreaming process that better addresses the needs and interests of all stakeholders. Effective communication and coordination among stakeholders, including local communities, national government agencies, and regional organizations, are also crucial for success. Although some strategies and approaches have been developed for mainstreaming biodiversity bringing together and addressing the needs of relevant stakeholders in the decision-making making process (Ginsburg et al. 2013; Redford et al. 2015; Whitehorn et al. 2019), so far relatively few have been implemented and have mostly remained conceptual ideas. Tailoring biodiversity information to user needs in the region can play a vital role in creating more effective policies for sustainable development that balance the needs of the environment and people.

In this study, we aimed to develop a codesign approach to mainstream biodiversity information in the Tropical Andes and create tailored biodiversity outputs that meet the needs of primary and secondary stakeholders in the region. A key intrinsic objective of the project was to establish networks and foster collaboration between stakeholders, including individuals, organizations, and countries involved. To achieve this, we employed a codesign approach that brought together key stakeholders and sectors that produce and use biodiversity information at local, national, and regional levels. We conducted surveys and workshops to identify priority needs and limitations in the flow of biodiversity information and to design indicators that address financial and technical capacity constraints. Additionally, capacity-building workshops were incorporated to improve the flow of biodiversity information in the region and address identified limitations.

## Methods

We developed tailored biodiversity products for stakeholders and sectors using a codesign process that was loosely adapted from a stakeholder engagement process outlined by Navarro et al. (2017). The process in this study was comprised of five steps: 1) engagement of stakeholders, 2) assessment of user needs and existing monitoring efforts, 3) codesign of biodiversity indicators, 4) implementation of biodiversity products, and 5) capacity building (Figure 2). To design and develop biodiversity indicators, we used the Essential Biodiversity Variables (EBVs) framework, which identifies a set of variables to monitor across genes, species, and ecosystems, as it enables the comparison of data across regions and sectors, identification of patterns and trends in biodiversity change, and provides a common framework for data collection, analysis, and interpretation (Pereira et al. 2013; Geijzendorffer et al. 2016; Pereira et al. 2017; Proença et al. 2017; Kissling et al. 2018).

**Figure 2.**
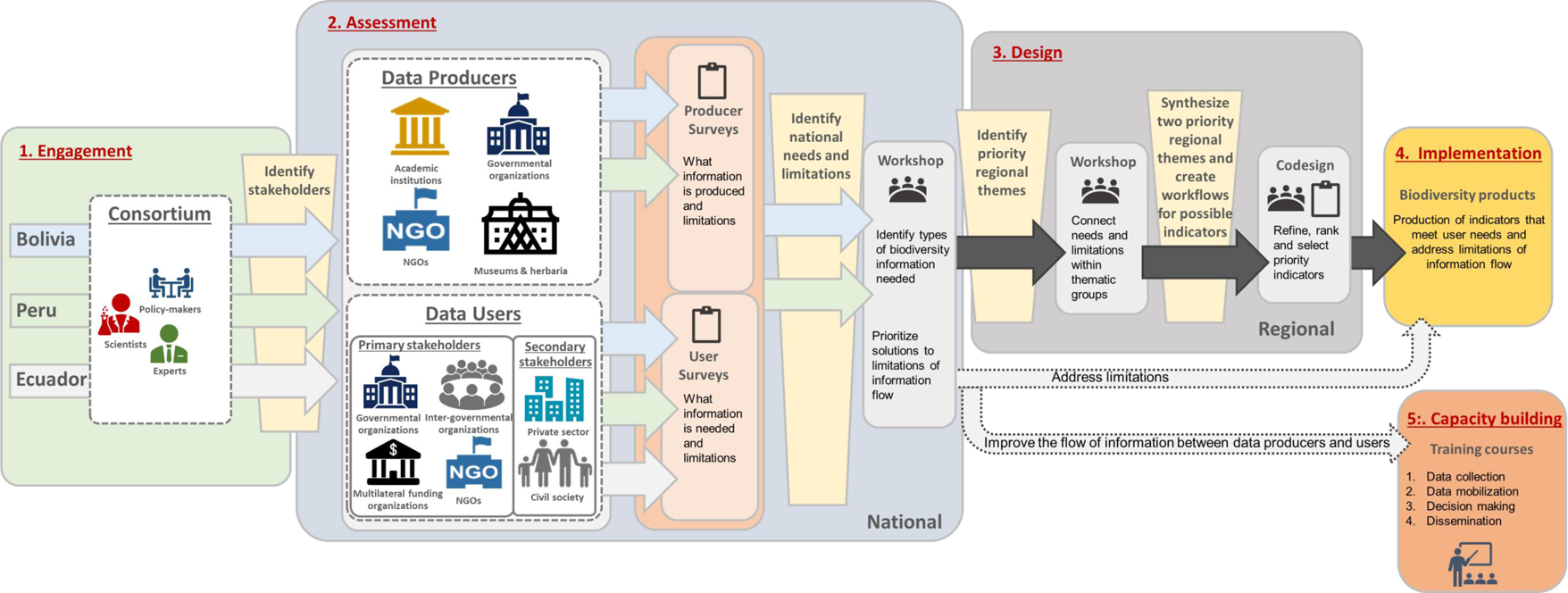
Flowchart depicting the codesign process used to identify priority needs and limitations of the flow of biodiversity information in the Tropical Andes region. The main outcomes of this process were the development of tailored biodiversity products that addressed major limitations and the improvement of information flow in the region through capacity building.

## 1 Engagement

For the first step, we engaged with scientific experts and policymakers within the region to establish a consortium of national partners and organizations to create an authorizing environment within Bolivia, Peru, and Ecuador (Figure 2). The institutions that participated in this initial process were the Instituto Nacional de Biodiversidad (INABIO) and the EcoCiencia Foundation in Ecuador; the Asociación Boliviana para la Investigación y Conservación de Ecosistemas Andino Amazónicos (ACEAA) in Bolivia; the Asociación para la Conservación de la Cuenca Amazónica - ACCA in Peru; as well as international partners including NatureServe in the USA, Universidad de Cordoba (UCO) in Spain, and the German Centre for Integrative Biodiversity Research (iDiv) in Germany. The engagement and knowledge of the national partners in the consortium were essential to identify key primary and secondary stakeholders of biodiversity information within their respective countries for the following steps of the co-design process.

## 2 Assessment

### 2.1 National Surveys

During the assessment phase, we sought to bridge the gap between biodiversity data producers and users in the Tropical Andes region by designing user-needs and data-producers surveys in collaboration with our regional partners (Figure 2). These surveys were sent to a diverse set of stakeholders, including decision-makers, scientists, NGOs, educators, citizens, and those in the private sector, and tailored to the linguistic and contextual variations of each country. Our objective was to identify and prioritize the needs of biodiversity data users, as well as gain a better understanding of the biodiversity data collection and management practices within each country.

We sent surveys to a total of 1,836 stakeholders for Peru and Bolivia, including 1,225 data producers and 611 users of biodiversity information. The survey process lasted for ten days, and email reminders were sent every three days to encourage responses. Although we were unable to survey data users in Ecuador due to COVID-19 financial constraints, we included a previous similar survey from 2018 that had been sent to 600 data producers in the country. In total, we received 443 responses (24.13% response rate) across the three countries. After integrating the outputs of the three national surveys, we identified the main limitations in the flow of biodiversity information and pinpointed priority thematic sectors specific to each country.

### 2.2 National workshops

The assessment phase continued with national workshops held by ACEAA and ACCA, aimed at refining the data needs identified in the surveys and identifying bottlenecks, as well as potential solutions to improve the flow of biodiversity information at the national level (Figure 2). Multiple stakeholders from different sectors were brought together in these workshops, providing a platform for users and producers to engage in dialogue and co-create mutually beneficial solutions. This allowed us to start identifying possible networks of people and institutions within each country who work on similar issues, who have the same information or knowledge needs, or where there could be a chain connection between information creators and users. Keynote presentations, forums, and group discussions were held in the workshops, gradually increasing the dialogue between different sectors involved in the production and application of biodiversity information. A total of 131 stakeholders participated in the workshops, representing academic, policy, societal, and economic sectors. Due to the COVID-19 pandemic, the workshops were held virtually to adhere to the restrictions on travel and in-person meetings. To address the multidisciplinary nature of the participants and communication barriers that exist between domains, the workshops used storylines and ecological narratives (Guerra et al. 2019) to ensure that all stakeholders spoke the same language and could efficiently communicate with each other. The results from the survey and workshops were distilled to prioritize six thematic groups common between the countries and fed into the regional workshop.

### 2.3 Regional workshop

To identify priorities for the Tropical Andes region regarding EBVs and the six thematic groups, a virtual regional workshop was held, inviting participants from the surveys and national workshops across the three countries (Figure 2). Due to the COVID-19 pandemic, the workshop was conducted virtually, with 131 stakeholders attending and an additional 188 listeners on YouTube and Facebook. The primary objective was to refine priority needs for biodiversity information that was specific to the six thematic groups distilled from the national workshops. To facilitate communication and prioritize regional needs, the workshop once again utilized storylines and ecological narratives, as described in the previous section. Participants were able to discover alternative approaches to producing, managing, developing, and utilizing information. They could exchange experiences and knowledge while finding opportunities for collaboration and synergy among individuals, organizations, and countries.

## 3 Design

To address limitations identified in previous workshops and meet the needs of stakeholders in the region, we engaged in a codesign process to develop user-relevant biodiversity indicators (Figure 2). We synthesized the six thematic groups into two priority regional themes and created a preliminary list of biodiversity indicators based on existing capacity and key spatial, temporal, and thematic priorities. We developed easy-to-understand workflows for each indicator to ensure accessibility and comprehension for all stakeholders. The coordination team then narrowed down the list to 8 indicators based on usefulness, validity, and feasibility. In a two-day co-design workshop, we collaborated with selected users, producers, and the consortium team to refine and develop two priority indicators for the Tropical Andes region. We presented an overview of the workflows and their connection to the regional themes, and stakeholders provided feedback on ways to enhance the indicators. We conducted a SWOT analysis on each updated indicator and discussed the results and their relevance and importance. The result was the selection of two biodiversity indicators that met the needs of stakeholders in the region. This co-design process allowed us to create indicators that were meaningful, feasible, and relevant to the region.

## 4 Implementation

The co-design process culminated in the production of biodiversity indicators (Figure 2) that were not only relevant to the needs of users but also scalable across different levels of governance. The indicators were designed to enable decision-makers to assess the status of biodiversity in the Tropical Andes region at different levels, from the local to the regional. However, the project also recognized that there were significant challenges in the flow of biodiversity data between data producers and users, which had to be addressed. To overcome these constraints, the project leveraged existing capacities and made the data accessible based on the FAIR principles (Findable, Accessible, Interoperable, and Reusable). This approach aimed to make the biodiversity indicators and its products more easily discoverable and accessible to stakeholders across the region, regardless of their level of expertise or location. By improving the flow of biodiversity information, decision-makers at different levels and sectors could access the information they needed to make informed decisions and contribute to the conservation and sustainable use of biodiversity in the Tropical Andes.

## 5 Capacity-building

To enhance the current and future flow of biodiversity information in the Tropical Andes region, a series of four capacity-building training workshops were conducted for stakeholders in data collection, analysis, decision-making, and dissemination (Table 1). The training workshops focused on building capacity in four key areas: 1) data collection and managing biodiversity information, 2) processing, analyzing, and synthesizing biodiversity information, 3) utilizing biodiversity information for decision-making, and 4) writing scientific papers and overcoming obstacles in the process (Table 1). The first three capacity-building workshops took place over three months and were attended by over 40 carefully selected individuals from 485 applicants from the Tropical Andes. The fourth and final workshop, a dissemination workshop, was held for 100 participants chosen from over 500 applicants beyond the three partner countries. This workshop included pre- and post-workshop surveys, which helped identify the main limitations and obstacles to biodiversity mainstreaming in the Tropical Andes and the topics they would like to see covered in future workshops. These workshops equipped the stakeholders with the knowledge and skills to better collect, manage, analyze, and disseminate biodiversity information, ultimately leading to improved decision-making and conservation efforts in the region.

**Table 1.**
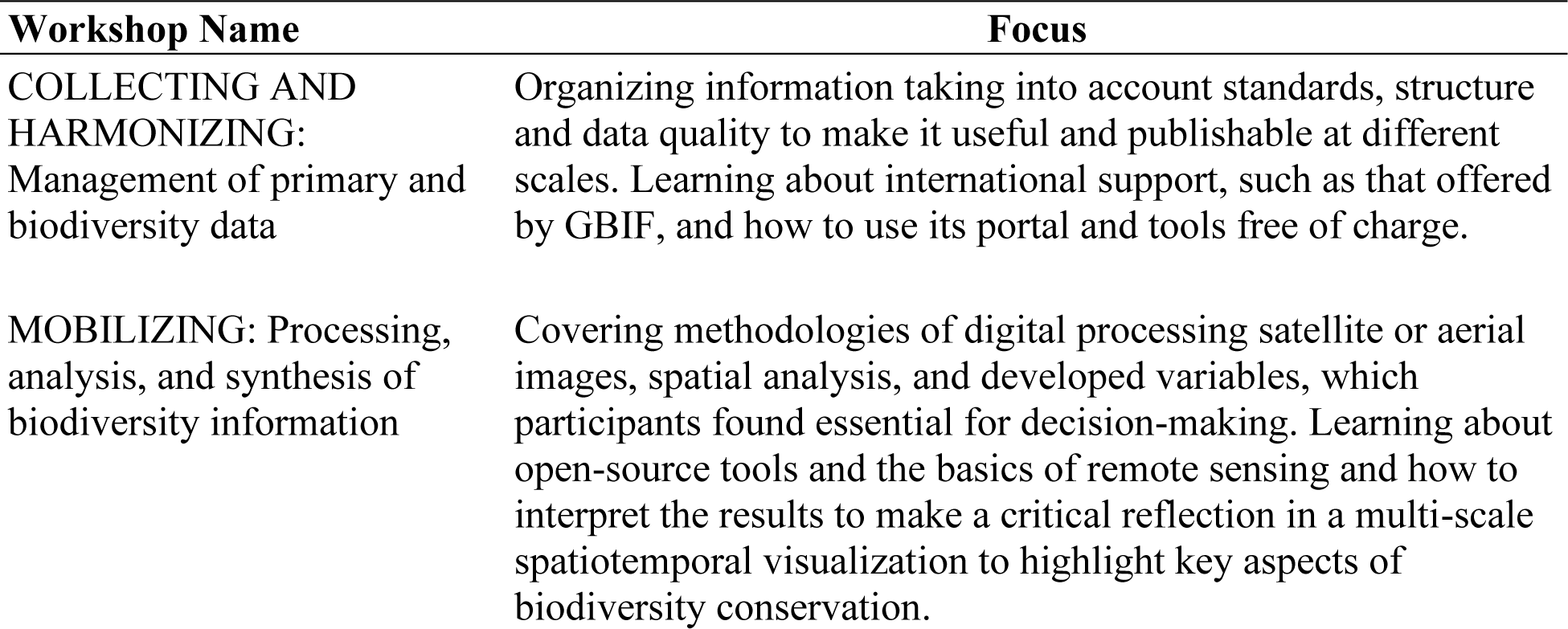
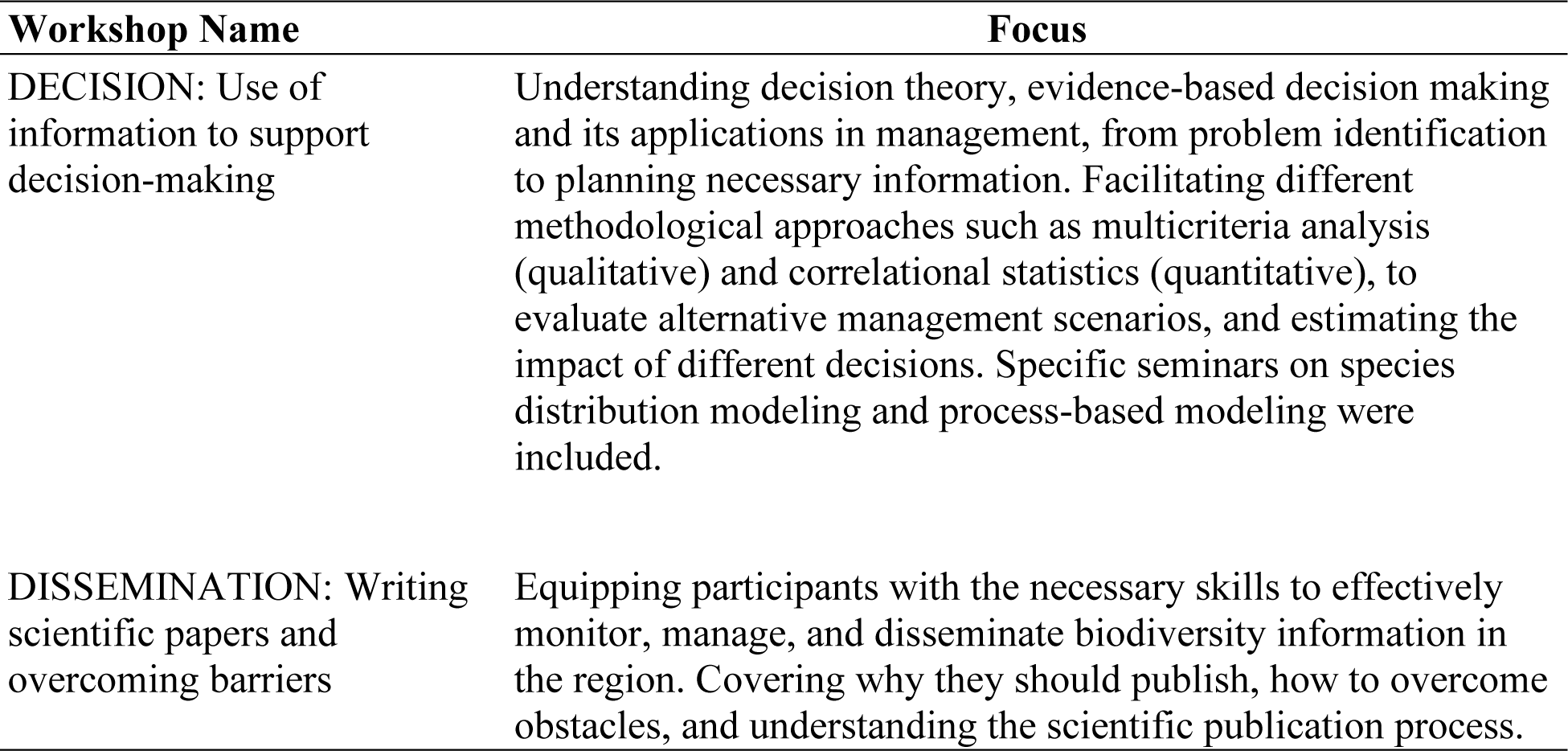
Summary of Biodiversity Capacity Building Workshops and Focus. The table shows the four modules of the workshop, their respective names, and the focus of each module.

## Results

### Stakeholders of biodiversity information

The stakeholder surveys revealed that the majority of those who identified as biodiversity data producers were researchers (83.94%), followed by non-academic executives/managers (9.76%). Users of biodiversity data were more diverse, including individuals from universities (25%), government (21%), private organizations (29%), and independent workers (21%). Among the users, the largest group was researchers (43%), with nearly equal affiliations between independent, government, and private institutions. Independent researchers, who work on short-term contracts or freelance, were commonly represented. Other secondary stakeholders included teachers (12.7%) and private sector professionals (11.07%). More than half (54.93%) of the survey respondents identified themselves as both producers and users of biodiversity data.

Obtaining valid and reliable data on the demographics and affiliations of participants in the national and regional workshops was a challenge due to the stochastic nature of attendance and participation. Nonetheless, we can confidently infer, based on our observations and the individuals we invited, that the majority of workshop participants belonged to three main categories: public citizens, private sector employees, and academic researchers from universities, research institutes, and NGOs. Another critical secondary stakeholder group was indigenous organizations, which participated in the workshops but were unable to respond to the survey due to the pandemic.

### Biodiversity focus and needs

The survey results revealed that stakeholders who utilized biodiversity information had a greater work emphasis on the social impacts of biodiversity in the region, while producers focused more on applied research. Both groups shared a general work focus on species, environmental management, and impacts, but there were notable differences between them (Figure 3a). Producers had a significantly higher specific focus on certain species aspects, particularly those related to endangered, migratory, and invasive species, as well as genetics and microbiology. While both producers and users had strong interests in environmental management (Figure 3a), producers had a higher percentage of work specifically focused on conservation management, while users had a stronger emphasis on natural resource management. Additionally, users had a more pronounced focus on general ecology as well as the social impacts of biodiversity, including economic factors such as food safety, tourism, and risk management (Figure 3a).

**Figure 3.**
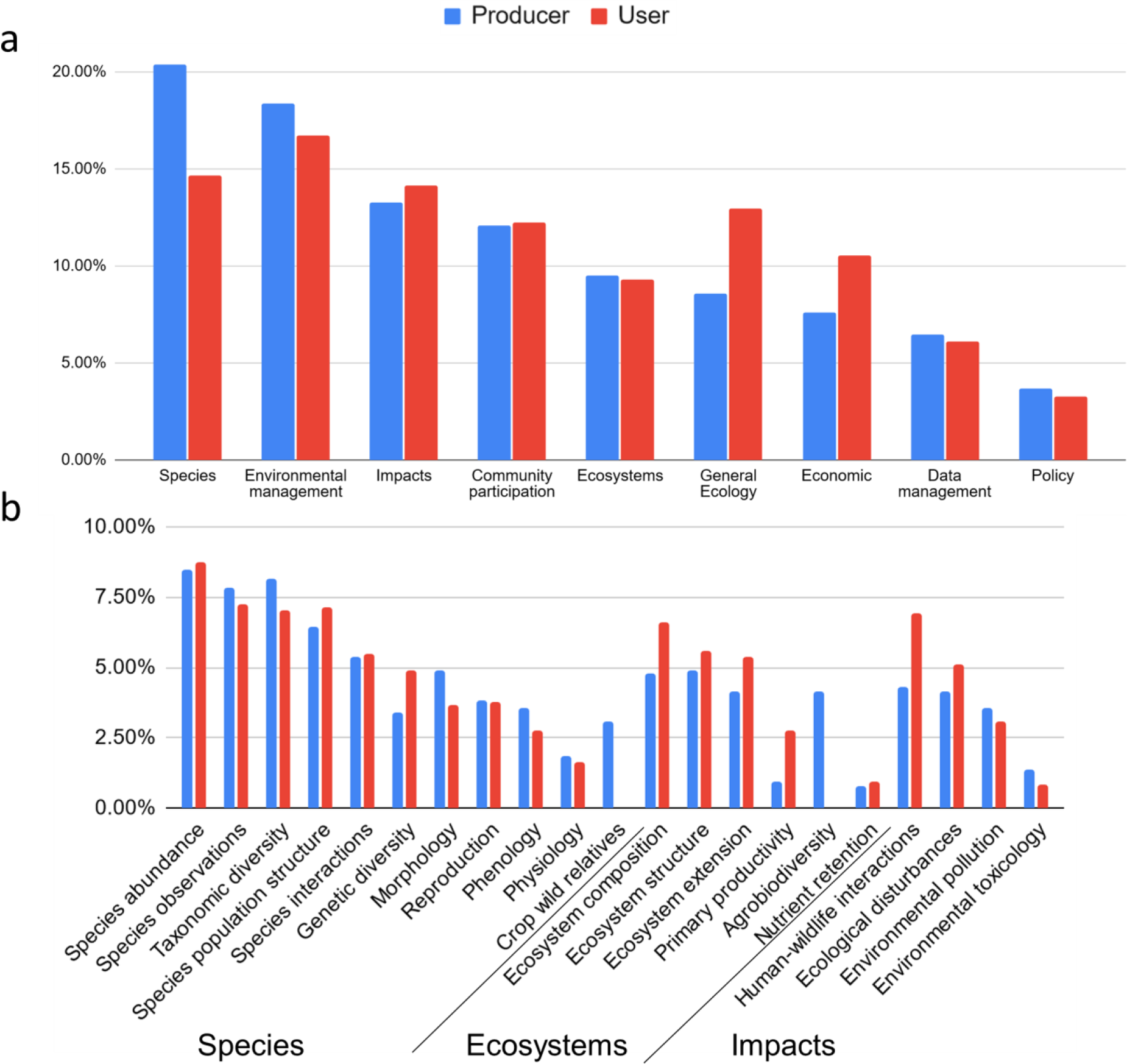
Percentage distribution of survey respondents who indicated a) the focus of their work involving biodiversity information and b) the types of biodiversity information they are interested in, according to producers and users in the Tropical Andes. The survey questions allowed for open-ended responses, and participants were able to select multiple options. The percentages represent the proportion of respondents who chose each option.

Based on the survey, the need for biodiversity information could be broadly categorized into three areas: species, ecosystems, and impacts. Species were the most frequently cited, specifically in terms of abundance, presence, and taxonomic diversity (Figure 3b). Ecosystem composition, structure, and extension (distribution) were the most commonly mentioned ecosystem topics, while human-wildlife interactions and ecological disturbances were the most frequently cited impact-related topics (Figure 3b).

### Limitations and bottlenecks

The survey results revealed that the highest-ranked limitations among stakeholders regarding the flow of biodiversity information in the Tropical Andes were bureaucracy, financial restrictions, and data accessibility (Figure 4). These limitations were further confirmed during the workshops. Funding limitations are particularly challenging due to the lack of resources available for biodiversity research globally, especially in underdeveloped countries with smaller GDPs like those in the region (Romero-Muñoz et al. 2019). This situation is further exacerbated in the Tropical Andes by national policies that prioritize development projects, infrastructure, and extractive industries over biodiversity concerns, as a response to socioeconomic challenges (Romero-Muñoz et al. 2019). The lack of funding, coupled with complicated bureaucracy, affects all stages of the information flow, from the collection of new information to updating, integration, technical management, and long-term storage and curation. Another obstacle lies in the absence of basic technological infrastructure and standardized guidelines for biodiversity information management, leading to fragmented knowledge and duplication of efforts. Establishing a robust technological infrastructure is an urgent requirement to improve efficiency and advance understanding and conservation efforts in the region.

**Figure 4.**
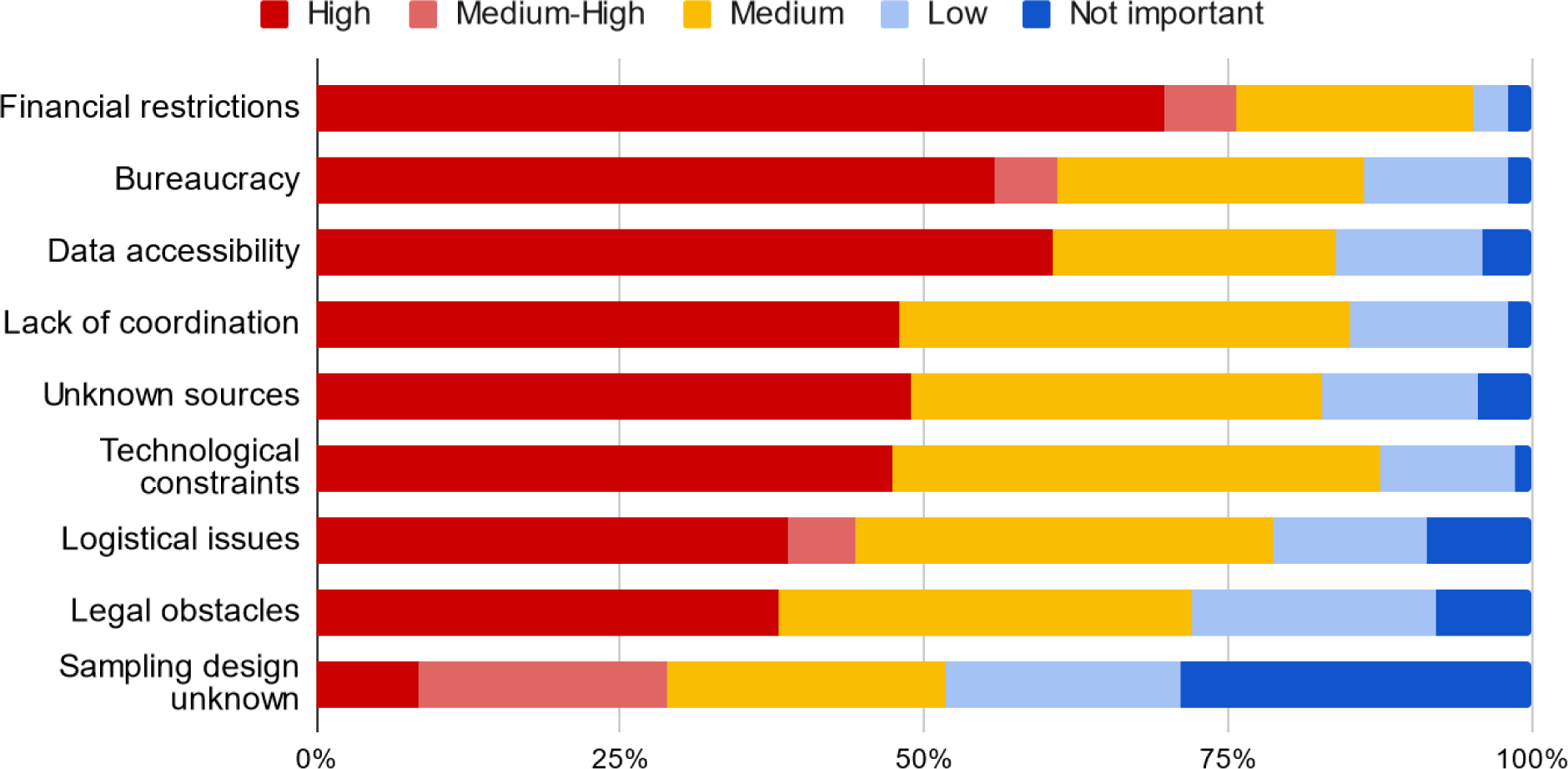
Main limitations and their importance for producing and obtaining biodiversity data in the Tropical Andes based on surveys.

Another major related limitation was the lack of accessibility to biodiversity information. Stakeholders reported that there is currently very little openly accessible data in the region, and many producers are either unwilling or unable to share their data openly. Technical capacity issues exacerbate this limitation, making overall access to information extremely limited. During the workshops, it was noted that many research articles were behind paywalls, even when the information was available, making much of the information unavailable to researchers, NGOs, indigenous organizations, and academic institutions that typically lack funds for these services in the region. Furthermore, researchers in Latin America typically do not receive competitive compensation for their publications compared to other regions (Owens 2022), which means many are not even able to access their own published research. Addressing these limitations is critical to improving the flow of biodiversity information in the region.

### Solutions to limitations and bottlenecks

During the national workshops, stakeholders proposed various mechanisms to address the aforementioned bottlenecks and facilitate collaboration and improve the flow of biodiversity information between data producers and users. The most popular solutions included making data and platforms freely accessible, ensuring publication and transparency of results, providing increased financial support, centralizing information and project results, developing cooperation networks, establishing clearer rules for data use, standardizing protocols, and using simple language and user-friendly platforms. To address these challenges, stakeholders suggested establishing transdisciplinary and inter-institutional spaces, such as stakeholder consultations, to facilitate collaboration between users and producers of biodiversity information and gain a better understanding of users’ needs. This would lead to more effective conservation and management strategies by creating a direct connection with relevant sectors of society and including them in the process of designing, implementing, and managing biodiversity information. Additionally, a network of knowledge exchange between scientists and stakeholders would address other suggestions such as creating cooperation networks, standardized protocols, using simple language, user-friendly platforms, and centralizing information and project results. This would also improve capacity building, which was identified as particularly important for the countries in this region. To address the limitation of publishing internationally, efforts were proposed to improve this through training and fostering international collaborations. Establishing a cooperation network would also allow for the other suggestions of making data and platforms more freely accessible, and ensuring publication and transparency of results. Overall, these solutions were incorporated, when possible, in the co-designing process of biodiversity indicators.

### Thematic groups

Based on surveys and national workshops, six thematic groups were identified as key priorities for biodiversity information flows: 1) industry, development, and infrastructure projects, 2) ecotourism, gastronomy, and national parks, 3) education and capacity building, 4) international mechanisms and agreements, 5) territorial planning and risk management, and 6) indigenous peoples and traditional knowledge holders (Table 2). These groups cover a wide range of activities and initiatives related to biodiversity conservation and management in the region.

**Table 2.** Thematic groups identified as key priorities for biodiversity information flows in the Tropical Andes region, as determined through surveys and stakeholder engagement in national workshops.

| **Sector** | **Description** |
| --- | --- |
| Industry, development, and infrastructure projects | Projects including mining, oil extraction, renewable energy, transportation, telecommunications, and pharmaceuticals. Agricultural insurance is also a growing sector in the region. |
| Ecotourism, gastronomy, and national parks | Activities related to scientific tourism, nature tourism, ecotourism, community tourism, and gastronomy in national parks. |
| Education and capacity building | Formal and informal educational initiatives related to biodiversity in the region, including capacity-building activities from primary school to university studies. |
| International mechanisms and agreements | Commitments made by the countries of the Tropical Andes region to comply with international mechanisms and agreements such as the UNFCCC Framework Convention on Climate Change, the UNCBD, NBSAPs, NDCs, and SDGs. |
| Territorial planning and risk management | Monitoring of both human and natural threats such as illegal deforestation, mercury in water, floods, landslides, fires, zoonotic diseases, and bio-prospecting pharmaceuticals, and collaboration between public and private health services. |
| Indigenous peoples and traditional knowledge holders | Utilization of traditional practices and knowledge to sustainably manage biodiversity through activities such as hunting, fishing, gathering, and agriculture, often in communal conservation areas. |

| Sector | Description |
| --- | --- |
|  | Also includes small-scale activities such as producing artisanal coffee, cocoa, and honey. |

### EBVs

The six sectors from the national workshops were used to map the requirements of specific EBVs (Appendix 1). Among the broader EBV classes required by all six groups, community composition, and ecosystem structure were identified as the most essential (Appendix 1Appendix). Specific EBVs were mainly community abundance, species abundance and distributions, and taxonomic diversity (Appendix 1Appendix). Genetic differentiation, inbreeding, and ecosystem phenology were not identified as relevant in any of the groups, while morphology and physiology, were discussed only in one thematic group (Appendix 1).

### Biodiversity indicators

We synthesized the six thematic groups into two regional priority themes: (1) land-use planning and risk management, which focused on large-scale development and infrastructure projects, and (2) international agreements and commitments, which emphasized the use of natural resources by local communities. Based on these themes and the relevant EBVs identified previously (Appendix 1), we developed 14 preliminary EBV-derived indicators that prioritized spatial and temporal considerations and the two priority regional themes (Appendix 2). The list was then narrowed down to eight indicators based on usefulness, validity, and feasibility (Figure 5a, with more details in Appendix 3).

**Figure 5.**
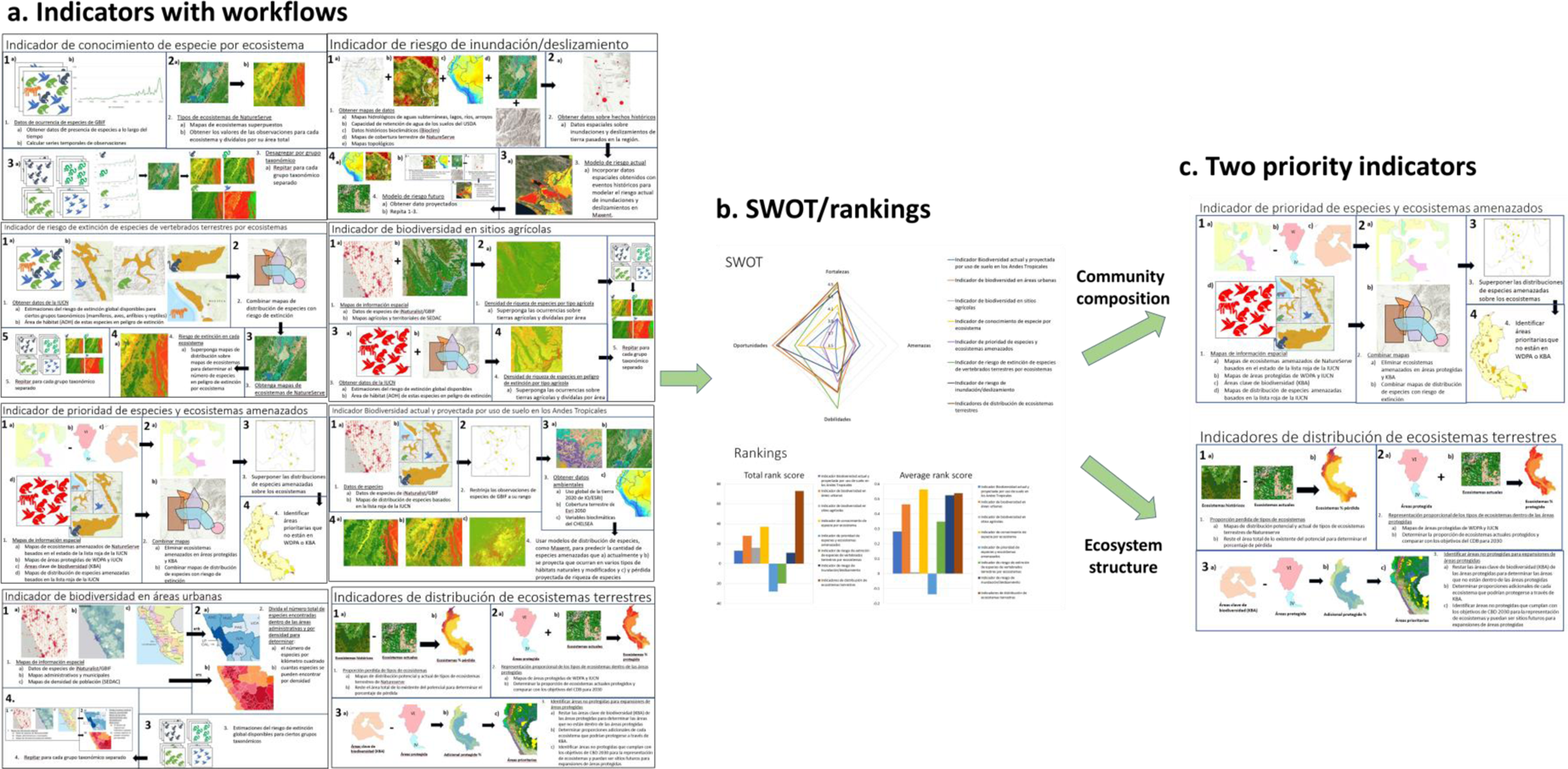
Workflows for the (a) eight biodiversity indicators that address the two priority regional themes in the Tropical Andes: land-use planning and risk management, and international agreements and commitments. The eight EBVs were then refined and prioritized (b) with SWOT analyses and rankings from a co-design workshop involving stakeholders. Based on this process, two EBVs related to “Community composition” and “Ecosystem structure” were selected that met the stakeholders’ needs in the region.

During the codesign workshop, we conducted an online SWOT analysis (Figure 5b) to select the most suitable candidates for the proof-of-concept design and implementation phase (Appendix 4). After further workshop discussions and final rankings (Appendix 4), we selected two indicators that met the stakeholders’ needs, derived from the “Community composition” and “Ecosystem structure” EBVs, namely “Species richness of terrestrial vertebrates by ecosystems” and “Terrestrial ecosystem distribution,” respectively (Figure 5c).

We utilized the NatureServe IVC hierarchical classification structure to develop our EBVs, allowing for scalable indicators that link measures of ecosystem diversity across different scales of conservation action (Comer et al. 2022). This hierarchical classification structure facilitates linking measures of ecosystem diversity across scales of conservation action. The regional assessment at the scale of the Tropical Andes provided insight into regional conservation and was readily scalable for reporting at continental or global scales. At the same time, these measures could be linked to ecosystem concepts defined and mapped for focused attention by land-use planners and managers working at more local scales.

The resulting EBV-derived indicators were instrumental in producing a highly impactful scientific paper (Comer et al. 2022). The paper presented a comprehensive regional analysis of the status and trends of biodiversity, as well as potential drivers in the Tropical Andes region. Moreover, the two biodiversity products were successful in addressing some of the limitations encountered during the assessment phase, such as the lack of openly accessible information that hindered the flow of biodiversity data between producers and users. For example, the manuscript and data available were developed based on the FAIR principles with the full geospatial datasets of the two scalable indicators openly accessible (Valdez 2023) and available for visualization in the GEO-BON EBV Data Portal (Valdez et al. 2022b, a). The manuscript also garnered significant media attention, with coverage in articles around the world and in different languages. A major article was even written in Spanish for Latin American audiences by Mongabay magazine, highlighting the broad impact of this work.

### Capacity-building

The workshops aimed to empower researchers and conservation practitioners in the Tropical Andes region, providing them with essential skills and knowledge for effective engagement in biodiversity conservation. The focus was on adopting a holistic approach to the information life cycle, emphasizing standardization and quality assessment of inventory data, management of international data publication platforms, utilization of open-source data and spatial analysis tools, and employing multicriteria analysis and correlational statistical modeling for decision-making. Initially planned as face-to-face sessions, the workshops were adapted to a virtual format due to COVID-19 restrictions. Despite the challenges, this transition facilitated participation from all three countries, allowing participants to exchange knowledge, connect with others who shared their concerns, gain insights into different realities, collaborate, and establish meaningful connections. Feedback from attendees of the virtual workshops was overwhelmingly positive, with participants expressing the value of the acquired skills and their applicability to their regular activities, as well as the potential for enhancing the quality of biodiversity information. Recognizing the benefits of ongoing communication and networking, participants strongly recommended the workshops to others seeking to improve their understanding and management of biodiversity information. The writing workshop was also highly rated by 93.5% of participants, who felt that the workshop exceeded their expectations. Pre-survey results for the writing workshop indicated that participants were interested in enhancing their abilities to convert their research findings into publishable papers, organize their information, and write effectively in English. Participants found the section on the publishing process to be the most valuable. They expressed a desire for future workshops to provide more detailed instructions on organizing information, navigating reviewer comments, and selecting appropriate journals for submission.

## Discussion

Our study aimed to improve the flow of biodiversity information in the Tropical Andes region by co-designing a process with a diverse group of stakeholders who produce and use this information. We identified priority biodiversity needs and current limitations, as well as bottlenecks and solutions to improve the flow of information. Our co-design process identified major limitations and incorporated commonly cited mechanisms to improve the flow of information between data producers and users. A key outcome was the co-design of two biodiversity indicators, which leveraged existing capacities while also addressing major limitations in the region. These indicators are easily linked to national land cover products in use, increasing their usability. We also incorporated commonly cited mechanisms to improve the flow of information, such as using simple language, freely accessible data, publication and transparency of results, establishing transdisciplinary spaces, using standardized protocols, and creating user-friendly platforms. Technical bottlenecks were tackled through capacity-building workshops for all stakeholders, which will facilitate the future flow of biodiversity information in the region.

Connecting stakeholders who produce and use biodiversity information was crucial for the codesign process, particularly secondary stakeholders such as local communities and economic sectors. Although this approach is not a novel concept (Kellert 1997; Díaz et al. 2015), it is often overlooked. These stakeholders play a key role in decision-making related to biodiversity conservation and management as they can impact and be impacted by biodiversity. Engaging secondary stakeholders can raise awareness and appreciation of biodiversity, leading to greater support for conservation. Local communities can also provide valuable knowledge, including traditional ecological knowledge, complementing scientific data and enhancing understanding of local biodiversity (Gewin 2022). Citizen science initiatives also further enrich our knowledge and lead to more informed decision-making. Incorporating diverse stakeholders fosters collaboration, builds trust, and promotes more effective and sustainable conservation efforts, considering diverse perspectives and needs. This is particularly important in regions such as the Tropical Andes, which may face “parachute” science, where research or programs are conducted without recognition of local governance, capacity, expertise, and social structures (de Vos & Schwartz 2022).

This study aimed to develop biodiversity information tailored to the needs of users in the Tropical Andes and test a model for mainstreaming biodiversity. To achieve effective mainstreaming, several actions are necessary, including engaging secondary stakeholders, facilitating the flow of biodiversity information from data producers to users, and incorporating the social and economic benefits of biodiversity into mainstreaming strategies (Figure 1). Scientists and policymakers should collaborate in participatory processes to ensure that biodiversity information is understandable and accessible to a broader range of stakeholders (Bickford et al. 2012; Davis et al. 2014). To achieve this, they can develop plain language summaries, use multimedia formats, and engage in targeted outreach and engagement (Novacek 2008; Diedrich et al. 2011; Bickford et al. 2012; Jolibert & Wesselink 2012). Additionally, efforts must be made to incorporate the social and economic benefits of biodiversity into mainstreaming strategies, which requires the development of clear policies and guidelines that balance the needs of different stakeholder groups (Smith et al. 2020; Xu et al. 2021). Translational ecology has recently emerged as an effective approach to integrating scientific knowledge into decision-making processes and making biodiversity information accessible to a wider range of stakeholders (Davis et al. 2014; Schwartz et al. 2017). Prioritizing bottom-up approaches that involve local communities in mainstreaming strategies can also ensure context-specific and responsive strategies that foster buy-in and ownership across the broader community (Diedrich et al. 2011; Pascual et al. 2021; Perino et al. 2021). By bringing together and engaging diverse groups, we can identify bottlenecks and determine ways to improve the flow of biodiversity information (Figure 1). Implementing these strategies can help us overcome the disconnect between academic research and the needs of biodiversity information of various stakeholder groups and achieve greater success in mainstreaming biodiversity considerations into decision-making processes across sectors.

The main suggestion to improve the limitations and facilitate the flow of biodiversity information in the Tropical Andes was to establish a transdisciplinary and inter-institutional biodiversity network. A sustained, user-driven, locally operated, harmonized, and scalable biodiversity observation network (BON), such as developed by the Group on Earth Observations Biodiversity Observation Network (GEO BON), could help achieve this by improving the acquisition, coordination, and delivery of relevant and timely biodiversity data to users (Scholes et al. 2012; Navarro et al. 2017; Walters & Scholes 2017; Kissling et al. 2018). Harmonized observation networks could optimize current observation efforts and data, and adopting an approach based on Essential Variables (EBV or EESV) could help identify biases and prioritize data mobilization and modeling efforts (Geijzendorffer et al. 2016; Navarro et al. 2018; Balvanera et al. 2022). Although the EBV-based indicator identified in this study provides only a snapshot of the current state of biodiversity and does not capture changes over time, it can serve as a valuable baseline for monitoring and detecting future biodiversity changes. Furthermore, it can also serve as a starting point for the development of an EBV for a BON in the region. More specifically, the production and subsequent use of these indicators can incentivize further, in-situ data collection to both verify and improve the accuracy in future iterations and form the backbone for a collaborative, trans-boundary monitoring approach for the region. The establishment of a Tropical Andes biodiversity network could consolidate data, improve discoverability, access, and utility of information, and serve as a valuable tool for monitoring and detecting changes in biodiversity. This approach has shown promising results in other regions, such as the Arctic, New South Wales in Australia, Colombia, and Europe with EuropaBON (Navarro et al. 2017; Moersberger et al. 2022; Pereira et al. 2022). The co-design approach implemented in this study and its outcomes can be used as a proof-of-concept of the BON development process that could be applied to other regions.

## Conclusion

Effective biodiversity conservation requires a collaborative and multinational approach that involves a diverse range of stakeholders, including local communities and the economic sector (Zador et al. 2015; Bravo et al. 2016). Achieving a balance between biodiversity conservation, and political, economic, and socio-cultural development requires effective integration and communication between scientific communities and organizations that use biodiversity information (Neßhöver et al. 2013; Cvitanovic et al. 2016; Pascual et al. 2021). To achieve this, producers of biodiversity information must directly address the needs of all relevant sectors of society when designing and implementing public policies and development plans (Huntley & Redford 2014; Redford et al. 2015). Managing priorities reciprocally can lead to better conservation of biodiversity while sustaining equitable use (Armenteras 2021). A bottom-up, results-based codesign approach that engages and considers the needs and perspectives of all groups that would benefit from biodiversity information can promote inclusive and responsive biodiversity mainstreaming and contribute to the successful implementation of biodiversity policies and conservation goals (Perino et al. 2021). Given the multiple worldviews, values, and knowledge systems between science, policy, and practice the process presented here can be a valuable blueprint to mainstream biodiversity information and make it more inclusive in the future.

## Acknowledgments

We thank all the local institutions in the Tropical Andes including Conservación Amazónica-ACCA, Asociacion Boliviana para la Investigacion de Ecosistemas Andino Amazonicos (ACEAA), Fundacion Ecociencia, Ecuador, Instituto Nacional de Biodiversidad de Ecuador (INABIO), and several international institutions including NatureServe, Universidad de Cordoba (UCO) in Spain, the Global Biodiversity Information Facility (GBIF), the German Centre for Integrative Biodiversity Research (iDiv), and the Group on Earth Observations—Biodiversity Observation Network (GEO BON) who worked together to document needs of biodiversity data users in the Tropical Andes. We acknowledge the support of the Environmental Ministry of Perú (specifically the Dirección de Diversidad Biológica), as they provided the large database of producers, the permission to use their logo in the invitations, and helped contact key institutions for the national workshop, which was particularly important in obtaining participants during the COVID-19 pandemic. We also thank ERANet-LAC for funding this project. Lastly, we also thank all the 2019–2022 workshop participants without whom the knowledge essential to this work could not have been generated.

**Appendix 1.**
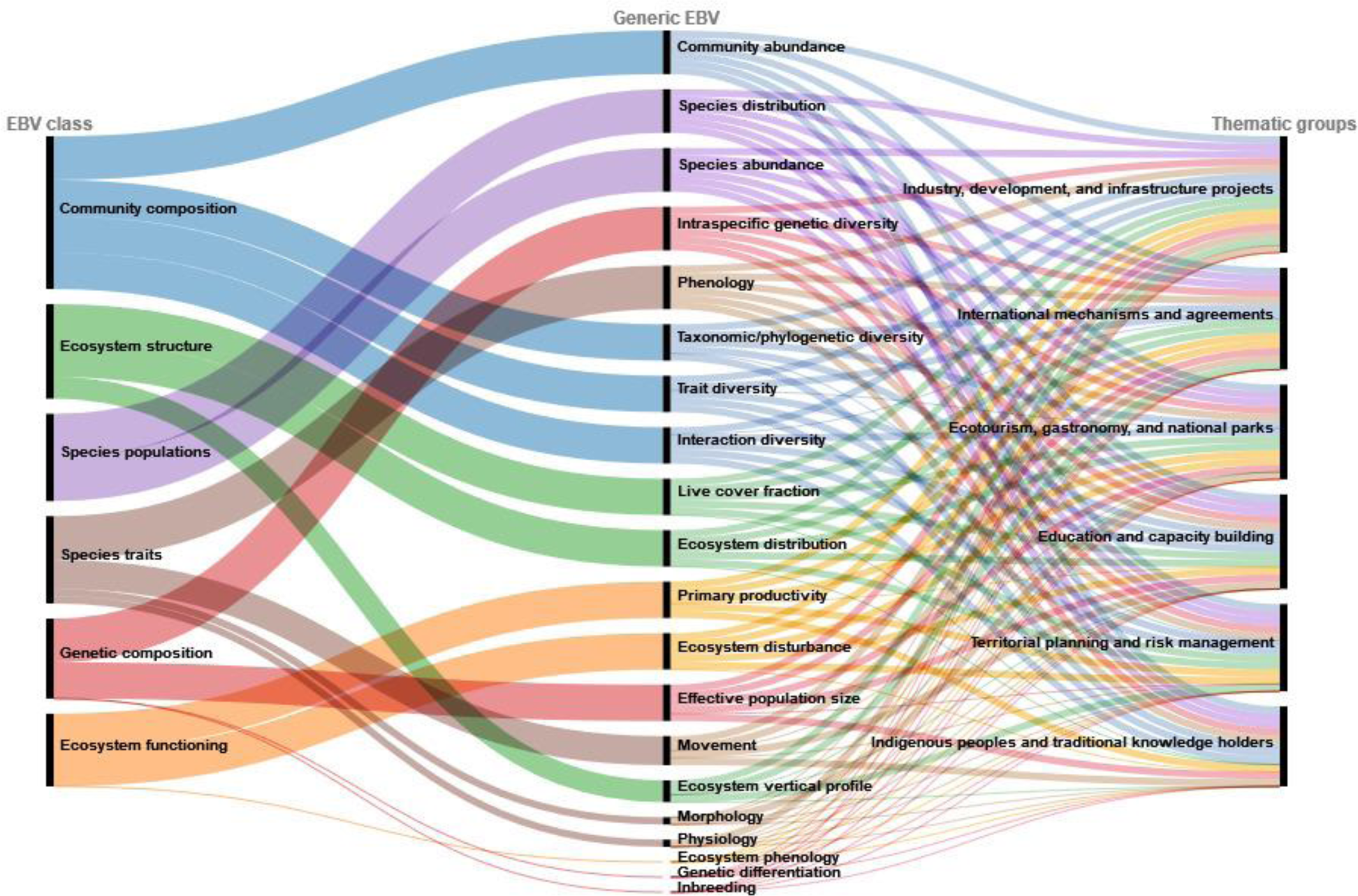
Priorities of Essential Biodiversity variables (EBV) for each of the six thematic groups identified in the Tropical Andes

Appendix 2. List of 14 preliminary EBV-derived indicators that drew on existing capacity and prioritized key spatial, temporal, and addressed two priority regional themes: (1) land-use planning and risk management, which focused on large development and infrastructure projects, and (2) international agreements and commitments. The first eight were the ones that were selected to be refined, revised, and ranked by stakeholders in the codesign workshop

### EBV based indicators for the Tropical Andes

**Objective:**

Provision of an initial list of EBV based indicators to be used in the Design Workshop of the Tropical Andes Observatory and that respond to the following priority topics:

- **T1:** Land-use planning and risk management focused on large development and infrastructure projects.
- **T2:** International agreements and commitments with emphasis on the use of natural resources by local communities.

Each indicator is graded A (Excellent), B (Good) to C (Fair) on three criteria:

<u>Useful:</u> is the EBV indicator beneficial, practical, and relevant in meeting the needs and supporting the ability of local partners to achieve and track conservation targets and sustainable development goals.

<u>Valid</u>: is the EBV indicator robust, well-grounded, and representative of what it is measuring.

<u>Feasible</u>: can the EBV indicator be effectively carried out and produce the desired result.

**1. Terrestrial ecosystems extent indicator (potential distribution, current distribution, current loss of potential habitat, representativeness in protected areas, additional representation in key biodiversity areas; across multiple levels of ecosystem classification)**

1. **Useful A**
2. **Valid A**
3. **Feasible A**

Topic addressed: T2

Documenting changes in ecosystem protection is essential to understanding the status of biodiversity and related ecosystem services. We will develop both potential and current distribution maps of terrestrial ecosystem types with the “potential” estimating where an ecosystem would likely occur today had there not been prior land conversion for modern land uses. We will assess the proportion lost and the representation of ecosystem types within protected areas as defined by IUCN I-VI protected area categories across multiple levels of terrestrial ecosystem classification based on the International Vegetation Classification (IVC) groupings. For ecosystem type representation within IUCN protected areas, they will be referenced to the potential extent of each type and compared with CBD 2030 targets for ecosystem representation. We will then calculate the additional proportions of each IVC ecosystem group that could be secured through identified Key Biodiversity Areas (KBAs). This will help us identify the protected areas with the most types of ecosystems for prioritization and unprotected areas with high numbers of ecosystems that can be future sites for protected area expansions. These indicators will help with CBD 2030 targets 1, 2, and 3.

Data required:

- Ecosystem maps (NatureServe)
- Protected areas categories (IUCN)
- World Database on Protected Areas (WDPA)
- Key biodiversity areas (KBA)

<u>Notes</u>:

**2. Species level knowledge indicator for each ecosystem**

- **Useful - A**
- **Valid - A**
- **Feasible - A**

Topic addressed: T1,T2

One of the main sources of information that provides a baseline of species-level knowledge is direct observations of species. This information on species identity and spatiotemporal reference traditionally comes from museums and herbaria, but has been recently supplemented with data from citizen science. We plan to analyze this species-level information in a way that helps us understand how much we know about a particular ecosystem. Using the most recent data from the Global Biodiversity Information Facility (GBIF) for each of the countries, we will analyze trends of information quantity over time for each ecosystem. This analysis will result in a time series composite indicator for all observations from the first year for which GBIF data is available to the most recent year, which will be aggregated by ecosystems. We will also be able to disaggregate this indicator into its components and obtain the same values (number of observations) by taxonomic group focusing and analyzing mammals, amphibians, reptiles, and birds separately. These indicators will help with CBD 2030 targets 1 and 3.

Datasets required:

- Global Biodiversity Information Facility (GBIF) data for the three countries
- Ecosystem level classification (NatureServe)

<u>Notes</u>: There’s been some thinking about this in light of the proposed Target 20 under the GBF and there might be some useful insights into how to disaggregate this data in different ways but also how to bring in different measures of knowledge into the indicator. This indicator can be useful to identify areas/ecosystems that are under-represented and could link to something similar to the approach that Colombia’s Biomodelos takes by guiding targeted efforts to collect species occurrence data. One issue is that the GBIF data will likely be changing relatively rapidly over time.

**3. Flood/landslide risk indicator**

1. **Useful A**
2. **Valid B**
3. **Feasible A**

Topic addressed: T1

The Tropical Andes is characterized as a significant risk for extreme weather events. Flooding is a major risk that will become more common and deadlier due to climate change and landslides have become a big issue in the region particularly during La Niña years. To protect its citizens, current infrastructure, and plan for future development projects it is necessary to predict and highlight the highest areas that have the largest likelihood to be affected by flooding and landslides due to extreme weather events, flash floods, La Niña, or glacial lake outbursts. To achieve this, we will use hydrological maps of the region as well as soil, land cover, and bioclimatic data from Worldclim to model the areas with the highest risk potential under various climate change scenarios. This will help prepare the countries within the Tropical Andes by identifying areas most at risk and to better evaluate future development plans, while also meeting target 1 of the CBD 2030 goals.

Data required:

- Hydrological maps
  - Groundwater
  - Lakes, rivers, streams
- Water Holding Capacity of Soils (USDA)
- Land cover maps
- Bioclimatic data
  - Past
  - Projected scenarios
- Deforestation maps

<u>Notes</u>: Feasibility depends on data. However, some data and analysis might be available as a baseline. Analysis has been done in certain parts of the region by the NASA SERVIR program, AmeriGEO efforts, and work done by Colombia’s Met Institute. We will check with Nancy Searby at NASA if they are pursuing this indicator. Otherwise, it is not clear how the modeling will work. Will also require known flooding events to validate the model.

**4. Terrestrial vertebrate species extinction risk indicator by ecosystems**

1. **Useful - A**
2. **Valid - B**
3. **Feasible - A**

Topic addressed: T1

Regardless of scale, geography, and activity, one of the direct effects of development and infrastructure projects is the increase in species extinction rates. Reducing this risk of species extinction is the main goal of most companies’ investment portfolios. We will therefore produce a spatially explicit indicator of the extinction risk in each ecosystem in the Tropical Andes region by combing the 1) estimates of global extinction risk available for certain taxonomic groups (mammals, birds, amphibians, and reptiles) as well as their area of habitat (AOH) generated by the red list of species, with the 2) ecosystem classifications available for the Tropical Andean countries. These indicators will help with CBD 2030 targets 1, 2, and 3.

Datasets required:

- Red list maps for vertebrates (IUCN)
- Area of habitat (IUCN)
- Ecosystem classification (NatureServe)

<u>Notes:</u> Taxonomic groups be disaggregated even further into finer groups where the data allows. Although a static indicator, it could also be updated annually. However, extinction risk has some serious lag-time response and may not be that great as a direct indicator for development impacts on extinction risk. Nevertheless, it is still useful for identifying areas that have already high pressure on vertebrates. It may also be insensitive to local actions unless you limit to range-restricted species.

**5. Biodiversity in urban areas indicator (species richness and threatened species)**

1. **Useful A**
2. **Valid B**
3. **Feasible A**

Topic addressed: T1, T2

Urbanization has and will continue to occur in the Andes. While urban development is a major threat and typically reduces biodiversity, it is also important to recognize that urban areas also harbor many species, particularly in parks and other outdoor recreational sites. Recognizing these species can not only lead to further protections of the species that coincide with human populations from future development but also help local communities appreciate and take pride in the nature around them. We will first take 2020 population density maps from SEDAC and divide them with municipality maps of the region to identify an average population density per municipality and department. We will then obtain GBIF data and subset these administrative areas to identify the total number of species found within them and divide by the size of the administrative areas to determine the number of species per square km, and also divide by the density of the population to determine how many species can be found per density. Lastly, we will also determine the cities and municipalities that have the most threatened species per sq km. This indicator will help address targets 1,4, and 12 of the 2030 CBD goals.

- Population density maps (SEDAC)
- Administrative and municipality maps
- iNaturalist/GBIF species data
- IUCN species data

<u>Notes:</u> Addressing and tracking trends in urban biodiversity may not be considered as important or of potential positive impact since damage within urban areas is (largely) already done. There will be a huge effort effect with municipalities near urban areas where scientists and other observers occur will have much more biodiversity data. We also need to account for species-area relationships and differing sizes of municipalities.

**6. Biodiversity in agricultural sites indicator (species richness and threatened biodiversity)**

1. **Useful A**
2. **Valid B**
3. **Feasible A**

Topic addressed: T1, T2

One of the major human impacts on biodiversity is due to agriculture. While habitat modification removes certain species from the environment, others may survive or even thrive in these new sites. It is therefore important to identify what types of agricultural habitats harbor high species richness and high levels of threatened species, to ensure these sites are managed sustainably while also involving local communities in the process. To do this we will look at GBIF data of all species as well as only threatened species across various agricultural land-use types. This will help policy and decision-makers to prioritize what agricultural lands they should focus on when involving local communities to better help manage biodiversity. The 2030 CBD targets emphasized are 1, 10, 14, 18, and 19.

Data required:

- Global agricultural land maps (SEDAC)
- GBIF species data

<u>Notes:</u> Mike has created indicators for Ghana and Uganda that looked at the impact of agriculture on ecosystems and protected areas, so the methodology is already established. Nevertheless, there is the issue of cause and effect: if an agricultural type has lots of threatened species, does that mean we need more of that agricultural type or less? Also, if an agricultural type is a major threat to biodiversity, the analysis will show few threatened species there. Will this be interpreted as a need for more sustainable ag practices or not? We also need to be careful of underlying biogeographical patterns of diversity since areas at lower elevations and with greater precipitation will have more species richness. Perhaps the measure is of agriculture types and matched local natural areas in the same life zone or large-scale ecological area? Additionally, we need to account for GBIF biases on where people go to make observations and collect data.

**7. Threatened ecosystems and species priority indicator**

1. **Useful A**
2. **Valid B**
3. **Feasible A**

Topic addressed: T2

Despite the large proportion of protected areas in the region, 72% of all species and 90% of threatened endemic species are still insufficiently covered with its protected areas no more representative of biodiversity than non-protected areas. Additionally, 77% of the protected areas in the Tropical Andes are located in areas that are less vulnerable to habitat change and exhibit low biodiversity irreplaceability, resulting in seeing conservation goals failing for more than half of all species in the region. To prioritize and expand protected areas it is important to identify what currently protected areas are the most important and what unprotected areas might be valuable to protect. For this indicator, we will look at threatened ecosystems and overlay the distributions of threatened species over protected areas to find sites with the most threatened areas. We will also look at threatened species and ecosystems that don’t overlap with or have less than a quarter of their range in protected areas, to find the areas that would result in more complete coverage of threatened species. These indicators are vital for targets 1, 3, and 4 of the CBD 2030 goals.

- Threatened species and their occurrence (IUCN)
- Ecosystem maps (NatureServe)
- Protected areas categories (IUCN)
- World Database on Protected Areas (WDPA)

<u>Notes:</u> This has linkages to the Ecosystem Protection Level methodology of the SBA. We also need to see how does it improve on or enhance the outputs of the CSIRO Protected Area Representativeness Indicator https://bipdashboard.natureserve.org/bip_metadata/protected-area-representativeness-index.html, which we also have on the TAO Dashboard? Perhaps better a measure is what percent of the countries’ threatened species occur in a protected area? Similar to the first indicator.

**8. Current and projected biodiversity by land use in the Tropical Andes indicator (species richness and threatened biodiversity)**

1. **Useful A**
2. **Valid B**
3. **Feasible C**

Topic addressed: T1, T2

Protecting biodiversity can only be achieved by first understanding how the current situation affects species distributions and forecasting the effect of future changes. As species are inherently linked to their habitat and environmental niche, we will use global land use data and bioclimatic variables and combine them with the occurrence of species from GBIF data and restricted within each species range. By using species distribution modeling, such as Maxent, we can then predict the number of species by taxa (birds, reptiles, amphibians, mammals) that currently occur in various natural and modified habitat types. Using future projections, we can then determine the changes in species richness for the next thirty years. This will also be done exclusively with threatened species to better determine which species will be the most threatened and which habitats should be of focus to protect from future changes. This indicator is important as it will be directly related to CBD 2030 targets 1, 3, 4, 9, 10, 12, and 14.

Data required:

- Global land-use 2020 (IO,ESRI)
- Esri Land Cover 2050
- Bioclimatic variables / CHELSEA
- GBIF species occurrence
- IUCN range maps

<u>Notes:</u> Similar to 10. Are the Esri LandCover 2050 projections credible? The model-based indicator outputs may not be valid when you go to projections. This is a one-off indicator and requires a massive investment in modeling.

**9. Environmental risk from agricultural and urban expansion indicator (threatened biodiversity and ecosystems)**

1. **Useful A**
2. **Valid C**
3. **Feasible A**

Topic addressed: T1, T2

One of the major threats to biodiversity is habitat loss due to urbanization. To assess the impact of future urban expansion in the region we will use the map of likely future areas of urban expansion up to the year 2030 (SEDAC) along with IUCN range maps of threatened species and ecosystems. We will identify areas with the highest risk of urban expansion and how it affects the future range of threatened species and ecosystems to determine the number of species that will have more than half their range reduced and those that will have nearly all of their range removed and thus likely to become extinct. This can be used to come up with proactive measures for future development. This will help with CBD 2030 targets 1, 14, 18, and 19.

Data required:

- Urban expansion up to 2030 (SEDAC)
- IUCN range maps
  - Species
  - Ecosystems

<u>Notes:</u> We can also use area of habitat maps. This assumes species don’t shift their ranges (e.g., due to climate change). This indicator can be updated over time.

**10. Biodiversity hotspots indicator (species richness, threatened species, and threatened ecosystems)**

- **Useful B**
- **Valid B**
- **Feasible A**

Topic addressed: T2

The Tropical Andes is one of the most biologically diverse areas in the world. However, to protect its species and habitats it is imperative to identify and prioritize hotspot areas. To do this, we will overlay IUCN range maps of species in the region to identify the number of species per 1km pixel using hotspot analyses. This will be done for specific taxa (birds, amphibians, mammals, reptiles), threatened species, and threatened ecosystems. By identifying these various hotspots policy and management can better meet their goals based on specific objections. These indicators will also help address targets 1, 3, and 4 of the CBD 2030 goals.

Data required:

- Threatened species and their occurrence (IUCN)
- Area of habitat maps (IUCN)
- Ecosystem maps (NatureServe)

<u>Notes:</u> It may be redundant or a subset of the criteria-driven analysis done for identifying KBAs. There will be a taxonomic bias toward tetrapods. Similar to the previous indicator.

**11. Ecotourism indicator for the value of biodiversity**

- **Useful B**
- **Valid B**
- **Feasible B**

Topic addressed: T1, T2

Ecotourism is a major source of income for many areas in the Tropical Andes, particularly bird watching of endemic and restricted species. However, it is not always clear what areas should be highlighted for ecotourism or protected from undergoing development. To demonstrate the value of biodiversity, the most popular ecotourism areas and their estimated number of visitors and economic impact will be determined by each respective country. The total number of endemic and restricted bird species per km squared area will be identified using GBIF and used to determine the economic value of bird species to each area. This indicator will demonstrate the most valuable tourist sites and tourist sites with a similar number of species that have the potential for greater economic impact. The result will help address CBD 2030 targets 3, 4,9, 12, 16, 20, and 21.

- Most popular ecotourism sites
- Total economic value of ecotourism
- GBIF bird data
- IUCN red list

<u>Notes:</u> There are many more factors that affect the value for tourism, especially ease of access and availability of infrastructure for tourism. Would this indicator further threaten remote areas to development? Additionally, we can also generate a list of species using Map of Life (MOL) for each PA in the region as they included this function for Peru, Ecuador, and Bolivia.

**12. Composite indicator for gaps in conservation of climatic zones in the Tropical Andes**

- **Useful B**
- **Valid B**
- **Feasible B**

Topic addressed: T1, T2

Although including biodiversity-related elements in land-use planning and risk management processes as EBV-based indicators strengthens this process, the real challenge is to cross domains and try to integrate information from various fields. In this sense, we propose to create an indicator that integrates EBVs with environmental information, specifically climate data. For this we propose to 1) use monthly climate time series (CHELSA) at a km squared and derive the principal components from monthly total precipitation and mean monthly temperatures, 2) specialize the main PCA orthogonal axis, 3) contrast these with the current system of protected areas and KBAs to define geographic zones with combinations of climate patterns that are not represented in area-based protection systems in the Tropical Andes, 4) as the next step these filtered zones will be bounded with NatureServe Ecosystem Classification to focus on forested ecosystems, and 5) in the last step using deforestation data we will define conservation priorities in climatic areas not represented in protected areas and KBAS where deforestation rates are relatively high.

Data required:

- CHELSA high-resolution climate time series.
- Ecosystem classification data (NatureServe)
- Protected areas data (WDPA)
- KBA data (KBA database)

<u>Notes:</u> We may want to bring in other data inputs beyond climate to analyze and guide how well Protected Areas cover different environmental envelopes. The SANBI Spatial Biodiversity Assessment approach is relatively similar and provides a simple indicator (Ecosystem Protection Level) that tracks progress in covering representativeness and areas that are highly threatened (using RLE classification).

**13. Environmental impacts of (illegal) mining indicator (threatened species and ecosystems)**

- **Useful A**
- **Valid C**
- **Feasible C**

Topic addressed: T1, T2

Mining is a major industry in the Tropical Andes as they are a major source of mercury, gold, and copper. However, much of this mining activity results in habitat destruction, pollution, and biodiversity loss. Additionally, illegal mercury mining has been a major threat to the local communities in the region from the pollution of watersheds. To protect threatened areas while also understanding the importance of mining as an economic necessity, it is important to identify not only areas with high mining potential but which will have the lowest impacts on biodiversity. To do this we will use a mineral resource assessment map from the USGS along with the mining threat index from SEDAC and overlay with IUCN range maps of threatened species and ecosystems. This will help us identify mining areas with a high potential of resources, such as gold and copper, that will have the lowest impact on threatened species and ecosystems. This will help meet CBD’s 2030 targets 1,3,4,7,14, and 15.

- Mineral resources maps (USGS)
- Mining threat index (SEDAC)
- IUCN range maps of
  - Threatened species
  - Threatened ecosystems

<u>Notes:</u> This approach is similar to the outputs from IBAT in that they produce and make available IUCN range maps to guide extraction projects towards lowering impact. We could also use AOH maps and consider using range-size rarity as a measure. Need to be careful with elevational diversity gradient since high elevation areas have fewer species, including fewer threatened species, and may therefore appear to be low risk for mining, but any activity could have a disproportionate effect on these high altitude species.

**14. Risk of Zoonotic emergencies indicator**

- **Useful - B**
- **Valid - C**
- **Feasible - A**

Topic addressed: T2

One of the consequences of the global Covid-19 pandemic, in addition to the direct effects on the health of the human population, is the effect on both public and private economies. The recent IPBES assessment of biodiversity and pandemics has revealed a direct link between the degradation of natural ecosystems and the probability of the emergence of new diseases of zoonotic origin. Increasing the frequency of contact in areas of high biodiversity increases the probability of a zoonotic jump to humans. Since the taxonomic groups with high levels of viral load are relatively well known: rodents, macro-and Microchiroptera, it is possible to create richness maps that overlap with deforestation maps to have an initial understanding of zoonotic risk. We will combine the 1) species richness maps for rodents, macro- and Microchiroptera with 2) deforestation data from Global Forest Change to estimate areas where the risk of new zoonotic jumps is very high.

Datasets required:

- Species distributions for mammals (IUCN Red List)

- Area of habitat maps
- Global Forest Change data

<u>Notes:</u> While there is an issue of high relevance with increasing understanding of the links between rapid land-use change and risk of zoonotic outbreaks, we have to be careful about correlating forest cover change with zoonotic risk as the events leading to spillovers involve many additional factors. We may want to look at the ASEAN Biodiversity Dashboard (go to biodiversity data tab and select zoonosis outbreaks) where data from WAHIS was used to track patterns in animal to human disease transmission events (cases, deaths, etc.). However, its value is a bit skeptical due to the lack of a measure of effort and the risk of misinterpretation or context on how this relates to biodiversity (again due to the multi-factorial nature of what leads to animal-human transmission). Additionally, primates may also be additional sources for zoonotic diseases. Overall, this is an interesting twist on T2 indicators as it assumes local community interaction with natural resources is a risk.

**Appendix 3.**
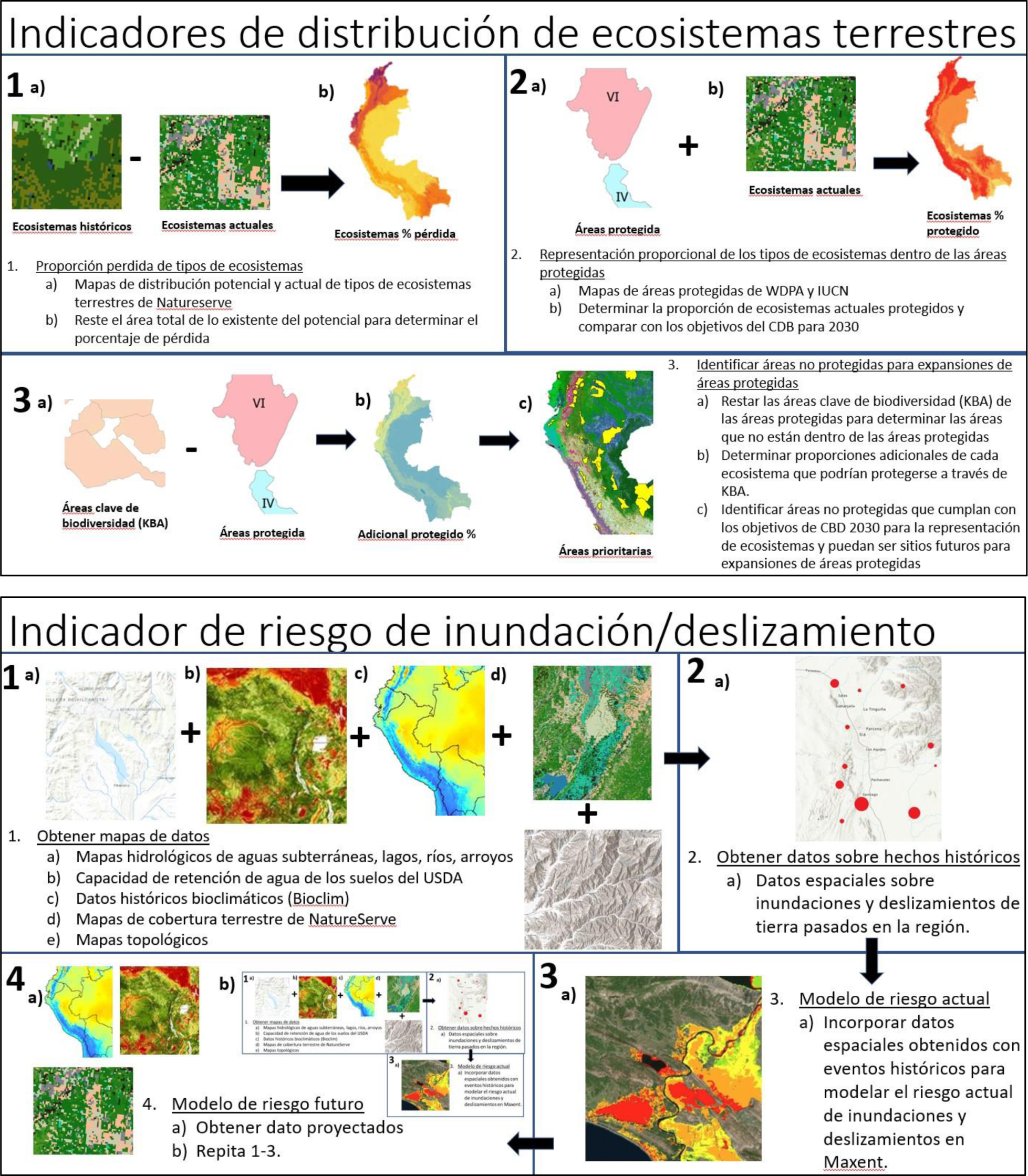
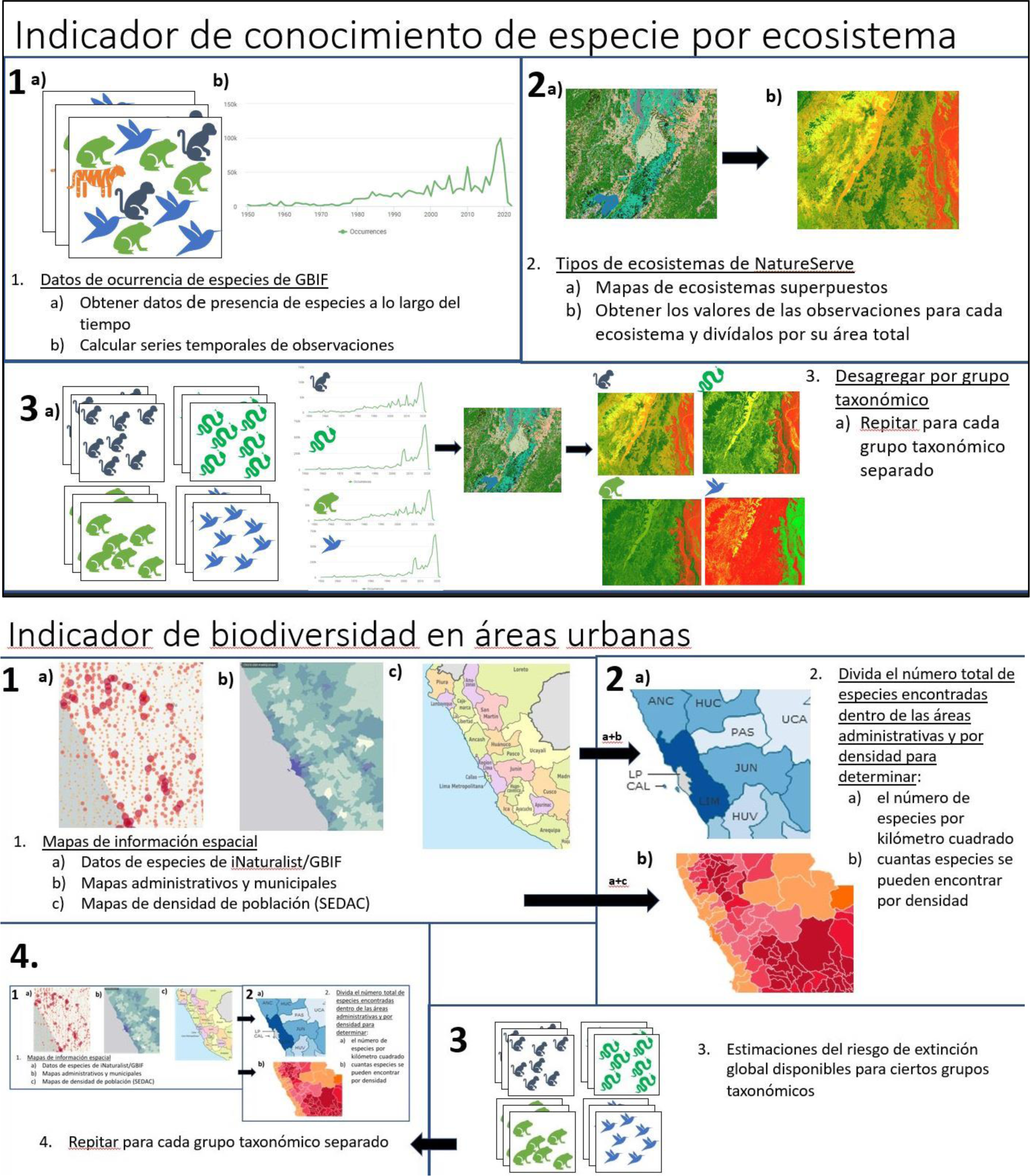
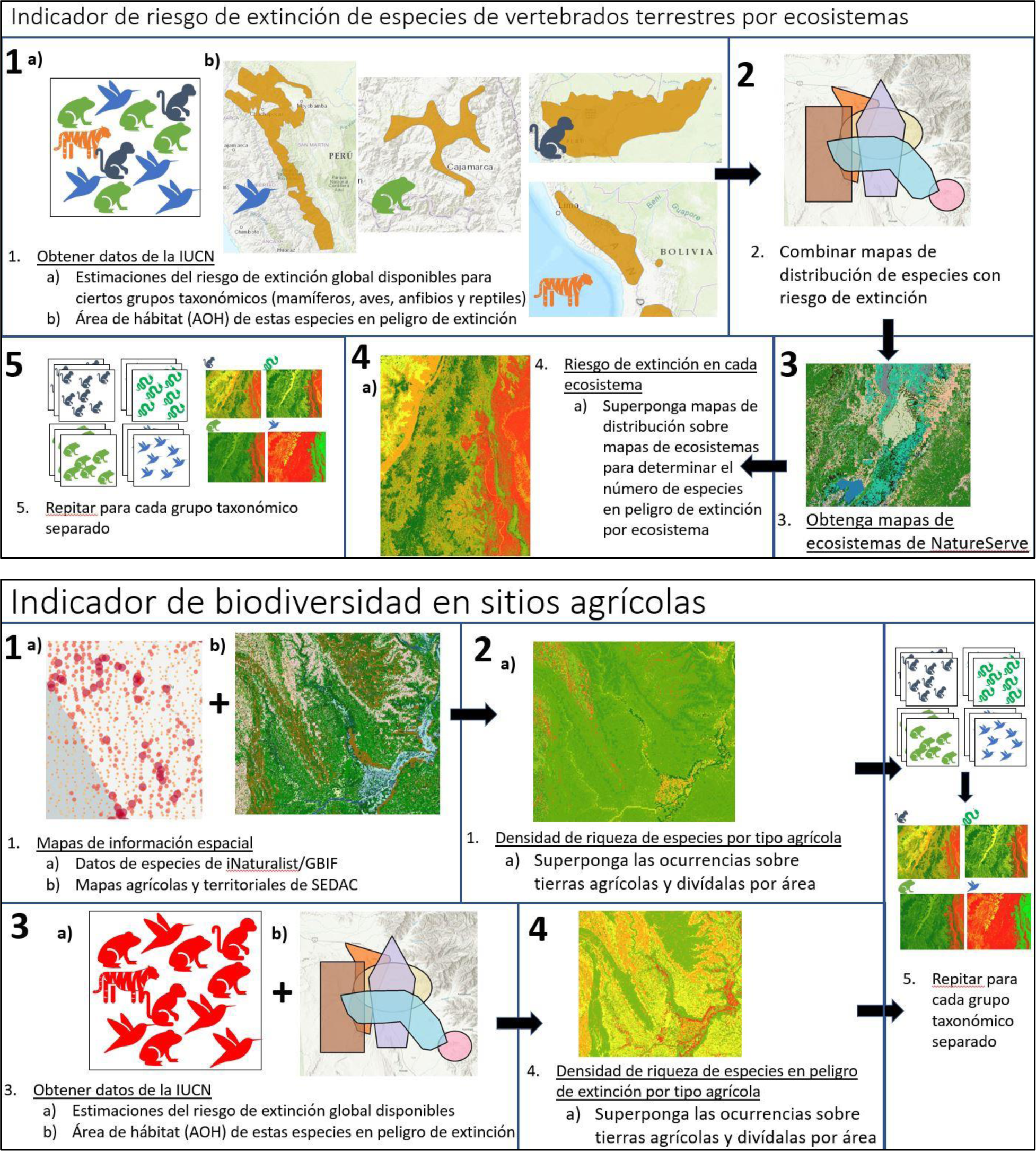
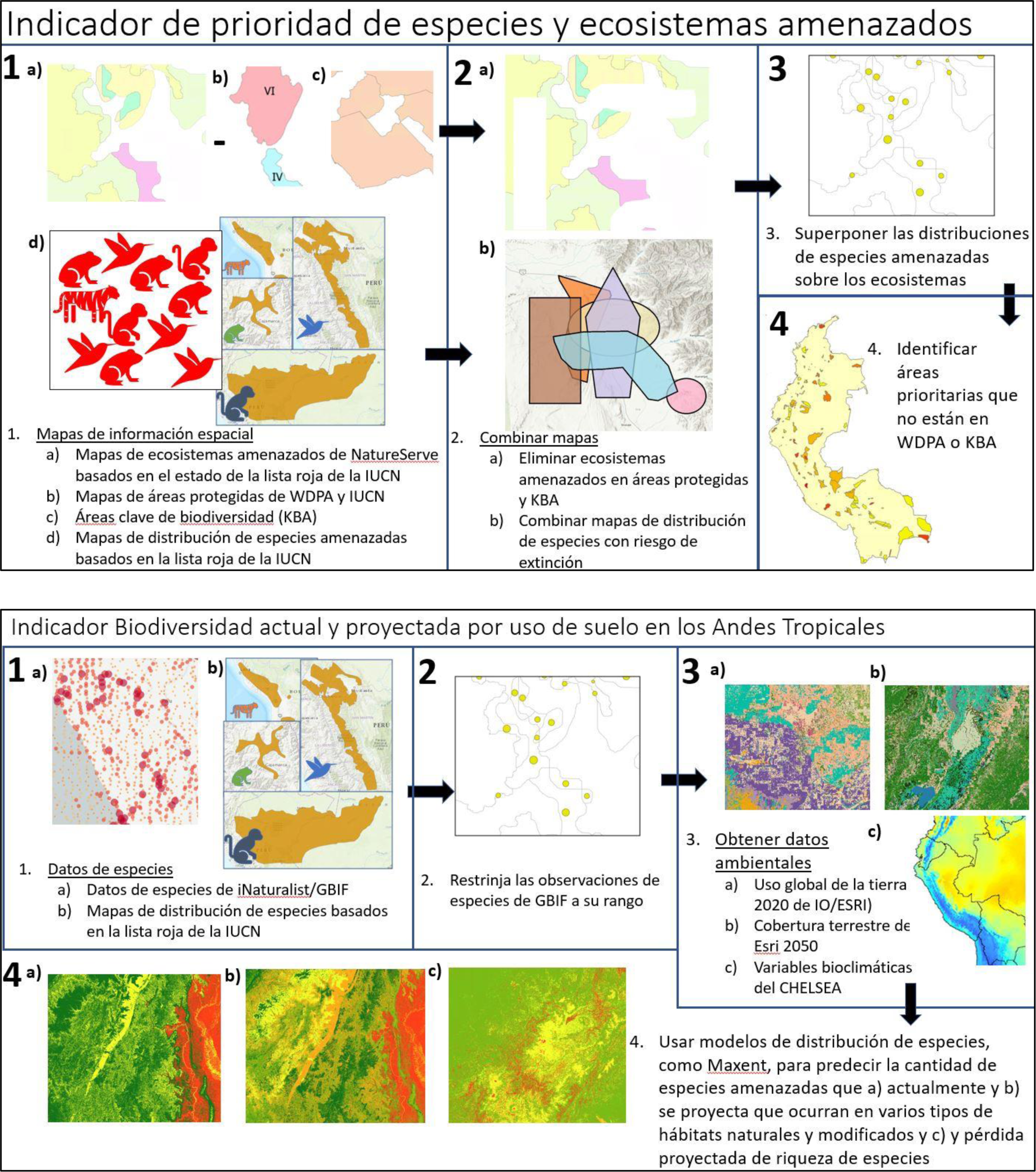
Workflows of 8 Essential Biodiversity Variable (EBV) indicators used in a codesign workshop to refine and select priority EBVs for the Tropical Andes region

**Appendix 4.**
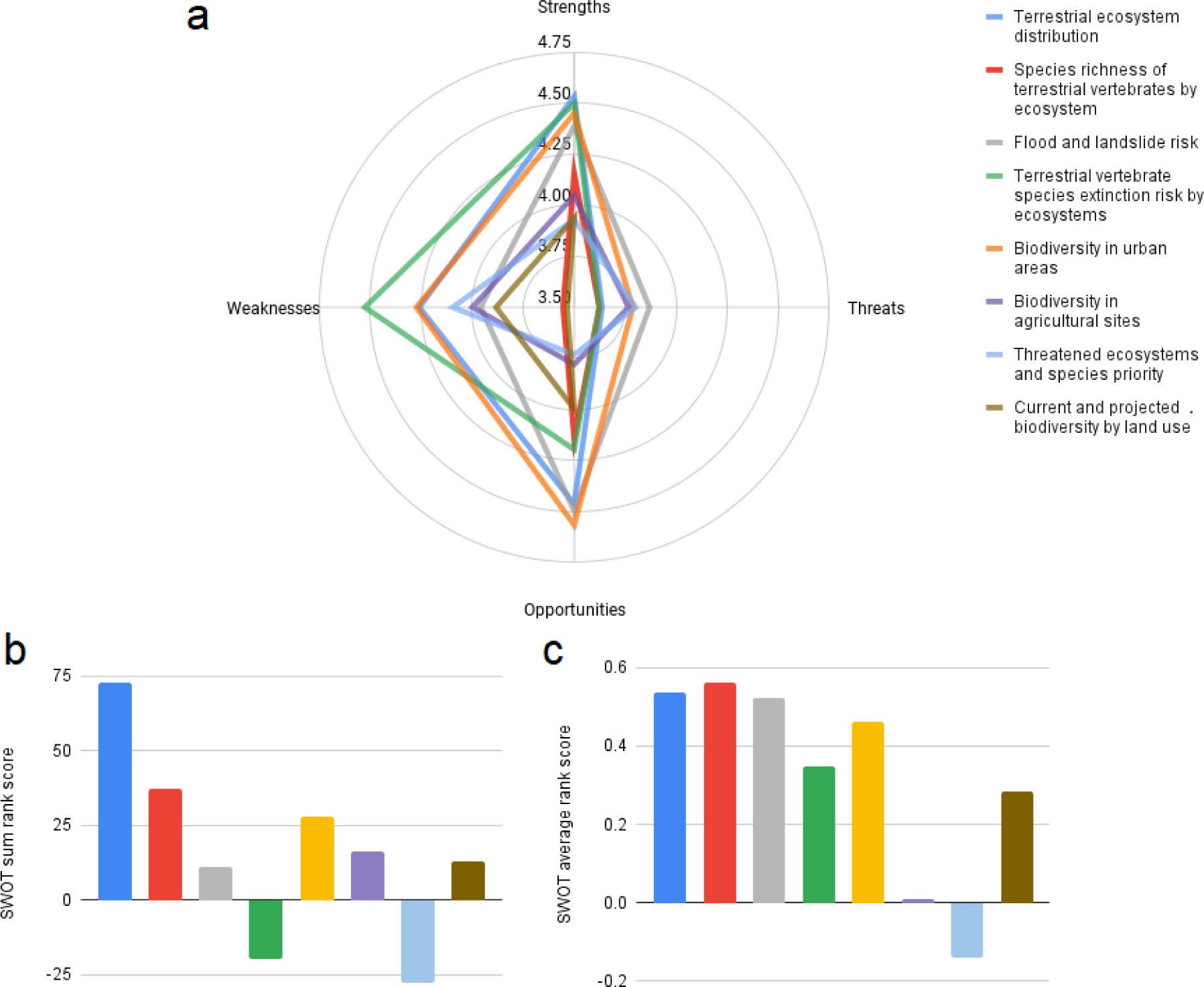
Appendix 4. Stakeholder SWOT (Strengths, Weaknesses, Opportunities, Threats) analyses from a codesign workshop of eight essential Biodiversity variables (EBVs) that addressed two priority regional themes in the Tropical Andes: land-use planning and risk management, and international agreements and commitments. The end result was (a) SWOT analyses with the average scores, (b) sum of scores (Sum of (S+O)-Sum of (W+T)), and (c) mean sum scores (mean S+ mean O)-mean W+mean T). Each participant provided various strengths, weaknesses, opportunities, and threats for each EBV and scored them on importance from 1(little important) and 5(very important)

